# Leveraging pleiotropy in genome-wide association studies in multiple traits with per trait interpretations

**DOI:** 10.1101/2020.05.17.100172

**Authors:** Kodi Taraszka, Noah Zaitlen, Eleazar Eskin

## Abstract

We introduce pleiotropic association test (PAT) for joint analysis of multiple traits using GWAS summary statistics. The method utilizes the decomposition of phenotypic covariation into genetic and environmental components to create a likelihood ratio test statistic for each genetic variant. Though PAT does not directly interpret which trait(s) drive the association, a per trait interpretation of the omnibus p-value is provided through an extension to the meta-analysis framework, m-values. In simulations, we show PAT controls the false positive rate, increases statistical power, and is robust to model misspecifications of genetic effect.

Additionally, simulations comparing PAT to two multi-trait methods, HIPO and MTAG show PAT having a 43.0% increase in the number of omnibus associations over the other methods. When these associations are interpreted on a per trait level using m-values, PAT has 52.2% more per trait interpretations with a 0.57% false positive assignment rate. When analyzing four traits from the UK Biobank, PAT identifies 22,095 novel associated variants. Through the m-values interpretation framework, the number of total per trait associations for two traits are almost tripled and are nearly doubled for another trait relative to the original single trait GWAS.

## 1 Introduction

Genome-wide association studies (GWAS) have been instrumental in identifying genetic variants associated with complex traits [Dorn and Cresci, 2009, Eskin, 2015, McCarthy et al., 2008]. As a result, there are tens of thousands of unique associations in the GWAS catalog [Milano et al., 2016]. With ever increasing sample sizes in GWAS more and more associated variants are being discovered. This suggests the presence of a large number of variants with small effect sizes that are not identified due to statistical power [Nishino et al., 2018]. As the number of traits examined as well as sample sizes increase over time, numerous variants are observed affecting more than one trait (i.e., pleiotropy) [Wang et al., 2010, Stearns, 2010, Gratten and Visscher, 2016, Chesmore et al., 2018, Visscher and Yang, 2016]. Some examples of pleiotropic effects include muscle mass and bone geometry, male pattern baldness and bone mineral density, as well as between multiple psychiatric disorders [Karasik and Kiel, 2010, Yap et al., 2018, Consortium, 2013].

We hypothesize that because variants often affect more than one trait, we can leverage this pleiotropy to jointly analyze multiple traits. This would potentially increase statistical power and identify variants with even weaker effect sizes. Following this intuition, there have been many approaches for performing association tests using summary statistics across multiple traits [Turley et al., 2018, Gai and Eskin, 2018, Zhu et al., 2015, Bolormaa et al., 2014, Qi and Chatterjee, 2018]. While simultaneously analyzing multiple traits is advantageous for identifying novel variants, performing an omnibus test is inherently difficult to interpret. This is because an omnibus test assigns one p-value per variant for the set of traits, and it is not clear how to assign a per trait significance level in this context. Even when this is done, it is not straightforward to interpret due to issues such as inflation in false discovery rates when the assumption of homogeneity in effect sizes is violated [Turley et al., 2018].

In this paper, we propose an alternative framework with a two step procedure. First, all traits are jointly analyzed to produce one p-value for each variant. If this p-value is significant, it suggests that the variant is associated with one or more of the traits. To accomplish this first step, we develop an efficient method called pleiotropic association test (PAT) which leverages the estimated genetic correlation between the traits to improve power and uses null simulations to accurately calibrate p-values. PAT also utilizes importance sampling to allow for estimation of significant p-values efficiently. The second step builds upon an interpretation framework first developed in the context of meta-analysis, m-values, to compute the posterior probability that a variant is associated with each trait [Han and Eskin, 2012]. We extend the m-values framework to take into account environmental and genetic correlation between traits.

In simulated data reflecting estimates of genetic and environmental covariance between real UK Biobank traits, we find that PAT is able to correctly control for false positives and increase power to identify novel associations. In comparisons to two multi-trait methods, MTAG and HIPO, PAT has a 43.0% increase in the number of associations over the next best method [Neale Lab, 2018, Qi and Chatterjee, 2018, Turley et al., 2018]. These results were then interpreted using the m-values framework where PAT identified 52.2% more per trait associations. While HIPO is shown to have similar power to MTAG for omnibus association testing, using the m-value framework to interpret HIPO’s associations resulted in a 40.3% increase in per trait associations relative to MTAG. We analyzed four traits in the UK Biobank where PAT identified 22,095 novel variants. We then use m-values to interpret the results for every trait. In two of the four traits, the number of per trait associations was almost three times greater than those found using the standard single trait GWAS, and it nearly doubled the number of per trait associations for another trait.

## 2 Results

### 2.1 Methods Overview

#### Pleiotropic Association Test

Here we introduce PAT (pleiotropic association test) which takes in GWAS summary statistics measured for T traits and assumes the z-scores follow a multivariate normal (MVN) distribution with mean 0 and covariance matrix *Σ*. Furthermore, it assumes the covariance matrix can be decomposed into two independent components, environment (*Σ_e_*) and genetics (*Σ_g_*). With this assumption in mind, PAT determines whether the genetic variant is associated with at least one of the *T* traits by performing hypothesis testing between two proposed MVN distributions using a likelihood ratio test (LRT). The null hypothesis is *Σ_g_* = 0; therefore there is only an environmental component. Under the null, the summary statistics for the variant *S* = {*s*_1_, …,*s_T_*} have the following distribution:

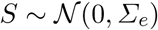

Under the alternative hypothesis, PAT models a genetic effect in every trait with effect sizes drawn according to the polygenic model and assumes the standard genetic correlation structure [Pasaniuc and Price, 2016, Rheenen et al., 2019, Aschard et al., 2017]. This results in there being both environmental and genetic components, so the z-scores have the following distribution:

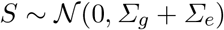

Having now defined the distributions, a LRT can be computed for each variant’s set of summary statistics S. Using the critical value *κ* for the threshold of significance, it can now be decided whether a variant is associated with the set of traits.

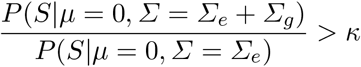

While likelihood ratio tests are known to approximately follow a mixture of χ^2^ distributions, PAT uses null simulations to set the critical value *κ* rather than a closed form solution. This is because utilizing a χ^2^ distributions can be complicated and may have reduced power [Visscher, 2006]. The critical value *κ* is set by simulating data under the null hypothesis *n* times, computing a likelihood ratio for each, sorting the set of likelihood ratios, and finally assigning a p-value to each likelihood ratio using the quantile. In order for the estimated critical value to be precise and reproducible, n needs to be very large. PAT is able to leverage importance sampling to compute critical values in fewer simulations (see Methods).

In simulations, PAT is compared to a version of standard GWAS generalized to multiple traits which we call multiple independent GWAS (MI GWAS). This approach is preferred to standard single trait GWAS as it takes into account multiple testing while being less stringent than a Bonferroni correction. MI GWAS determines if the genetic variant is associated with at least one trait if the largest |*s_i_*| ∈ *S* = {|*s_1_*|,…, |*s_T_*|} > *c*.

#### Multi-Trait GWAS Interpretation

While PAT is a powerful tool for determining if a genetic variant is associated with a set of traits, it provides only one p-value per variant because it is an omnibus test. This means that while it may reject *Σ_g_* = 0, there is no information as to which trait is driving the association. Therefore, we propose a means of interpreting the p-value on a per trait level by estimating the posterior probability of a variant having a non-zero effect on a trait. This framework, m-values, was originally developed for interpreting meta-analysis across studies, but here it is extended to account for the covariance structure between traits [Han and Eskin, 2012].

To provide some intuition on m-values, we will describe the P-M plot (p-value by m-value plot) [Han and Eskin, 2012] in Fig. 1. This plot has the per trait p-value from the original single trait GWAS along the y-axis and the per trait m-value along the x-axis. A line at − *log*(5 × 10^−8^) denotes the threshold where a variant is considered genome-wide significant. Region A is where the original single trait GWAS resulted in the variant being significant while the interpretation of the omnibus test did not. There should be no data points in this region.

**Fig. 1:**
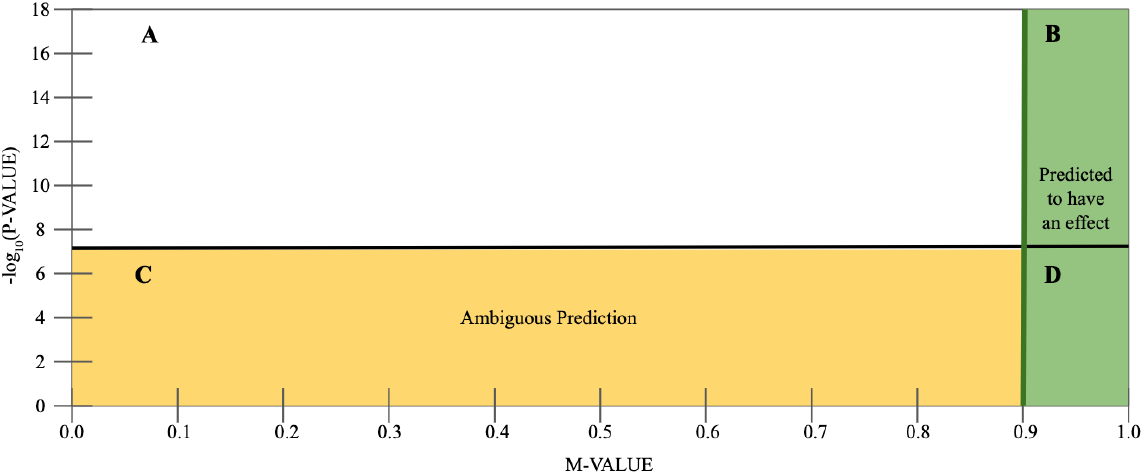
A figure depicting m-value interpretations using a P-M plot. Along the x-axis is the per trait m-value while along the y-axis is the p-value from the original single trait GWAS. Region A is when the original association is significant, but the m-value interpretation is ambiguous. There should not be data points in this region. Region B and D are associations with an m-value greater than 0.9, so the interpretation is there is a genetic effect in this trait. In Region C, the m-value interpretation is left ambiguous.

Regions B and D contain variants interpreted as associated with the trait because the m-value is greater than 0.9. Some of these variants have already been identified by the single trait GWAS (B) while other traits will be uniquely discovered on a per trait level (D). Region C contains the variants whose m-value is less than or equal to 0.9 and were left with an ambiguous interpretation.

### 2.2 Covariance structure between traits impacts the shape of PAT’s rejection region

In order to provide an intuition on and compare the rejection regions for MI GWAS and PAT, we simulate 100,000 summary statistics for two traits and set the genetic variance (*σ_g_*) equal to 0.0007 for each variant in both traits. The level of significance is set to *α* = 0.05, and the results are shown in Fig. 2. The first column highlights MI GWAS’s performance. Variants which were correctly identified as associated are shown in red while the ones missed by MI GWAS are grey. In each row regardless of model specification, the shape of MI GWAS’s rejection region is a square. As MI GWAS does not account for genetic correlation, there is no effect on the critical value when this parameter varies.

**Fig. 2:**
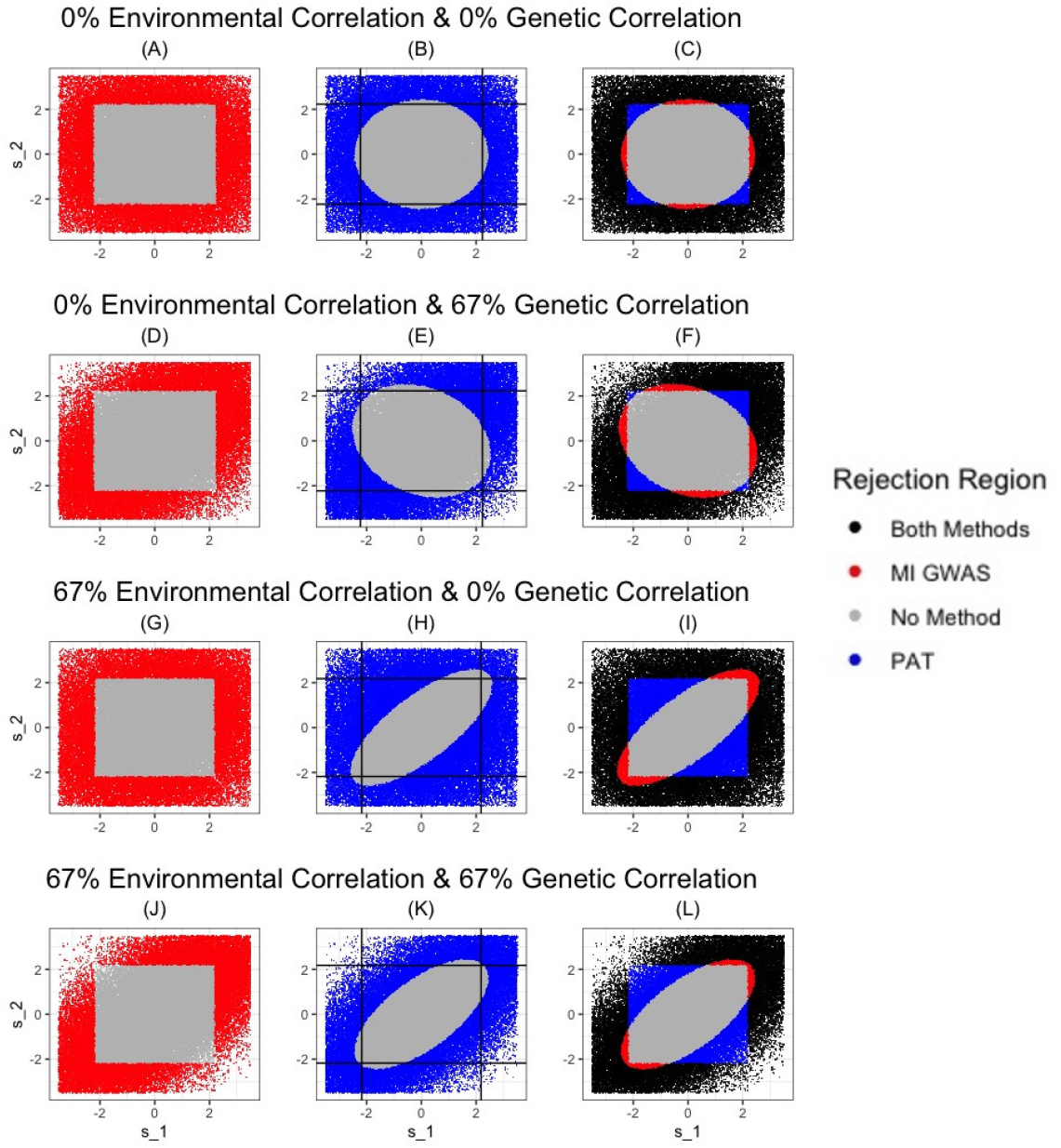
Comparison of the rejection regions for MI GWAS and PAT. We simulate 100,000 summary statistics for two traits and set the genetic variance *σ_g_* = 0.0007 for both traits. We vary the genetic and environmental correlations and use *α* = 0.05 level of significance for comparing the methods. Each row of figures is one set of simulations highlighting 3 points. The left column shows the rejection region of MI GWAS, the middle column is PAT’s rejection region while the third column is a comparison of the two methods. The simulations used in sub-figures A-C have no environmental or genetic correlation while the data in sub-figures D-F have no environmental correlation and 67% genetic correlation. For the third row of sub-figures, G-I, the environmental correlation is 67% while there is no genetic correlation between traits. The last row of simulations assume an environmental and genetic correlation of 67%.

The same phenomenon is not true for PAT which is shown in the second column (with the critical values of MI GWAS depicted with black lines). Here the data points for which PAT rejects the null hypothesis are shown in blue, and those PAT failed to reject are depicted as grey. In all four rows the shape of the rejection region is an ellipse. As PAT models environmental and genetic correlation, both parameters impact the shape of the elliptical rejection region. In the first row there is no environmental or genetic correlation, so the shape is exactly a circle. This means any extreme value for at least one of the summary statistics is likely to be rejected. In the second row, we model 67% genetic correlation and no environmental correlation. Here, the shape enables PAT to correctly identify more variants with positively correlated z-scores but fails to aid in identifying variants with negatively correlated z-scores. This follows the intuition that modeling genetic correlation would increase power to identify variants whose summary statistics follow this correlation pattern.

While the first two rows follow intuition, the shape of PAT’s rejection region in the last two rows is less intuitive. In the third row, we simulate traits with 67% environmental correlation but no genetic correlation. In this situation, the shape of the rejection region is in the direction of the environmental correlation; therefore, PAT has more power when z-scores are negatively correlated relative to when they are positively correlated. This means when summary statistics are positively correlated, PAT fails to reject the null unless the values are very extreme because it assumes the only source of positive correlation is the environment. The gain in power in the direction of negative correlation is due to the same idea that these values are unlikely under a positively correlated environment unless there is a non-environmental effect (i.e., genetics). In the final row, we simulate with a positively correlated environment and genetics. Here, the shape of the rejection region still follows the direction of the environmental correlation. This aids in controlling false positives, but it means that PAT may also be overly conservative in the direction of environmental correlation even when there is genetic correlation in the same direction. Further to that point, the critical value for MI GWAS as shown in sub-figures H and K (black lines) is identical. In sub-figure K, there are fewer variants pass this cut-off that are missed by PAT relative to sub-figure H. This mean that while PAT will be always be more conservative in the direction of the environmental correlation, it will be less conservative when it actually expects a genetic reason for correlated summary statistics.

The right most column is a comparison of the relative power of PAT and MI GWAS. Variants that were correctly identified by both methods can be seen as black data points while those missed by both are shown in grey. The variants only identified as significant by PAT are blue while those found only by MI GWAS are red. Under all four simulation frameworks, PAT has more statistical power than MI GWAS with the greatest improvement occurring when there was genetic correlation (sub-figures F and L). When there is no environmental correlation, the improvement is largest. We note that the size of the blue region may appear smaller in sub-figure F than in sub-figures I and L, but the density of the data points is higher due to the power increase being in the direction of genetic correlation.

### 2.3 PAT controls false positives

We select four quantitative traits: body mass index 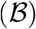, diastolic blood pressure 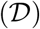, height 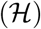, and systolic blood pressure 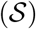 from the UK Biobank to simulate z-scores reflective of real data (see Appendix) [Neale Lab, 2017]. We simulate 10^8^ summary statistics with 25 replications according to the environmental correlation estimated from the UK Biobank. While the simulations only have environmental correlation, PAT also models the alternative hypothesis which assumes a genetic effect and genetic correlation between the traits (see Methods). Here, the calibration of MI GWAS is also checked, but this method makes no assumptions about the phenotypic covariance between traits. Fig. 3 shows box-plots of the proportion of p-values below a significance threshold *α* for PAT and MI GWAS. The threshold *α* is set to the following values: 5%, 2%, 1%, and 0.5%. As the distribution of p-values under the null hypothesis is uniform, 5% of p-values are expected to be smaller than the level of significance, *α* = 5%. This expectation holds for all levels of significance under the null. In Fig. 3, both methods are shown behaving within expectation at the various levels of significance. This indicates both methods are effective at controlling the false positive rate.

**Fig. 3:**
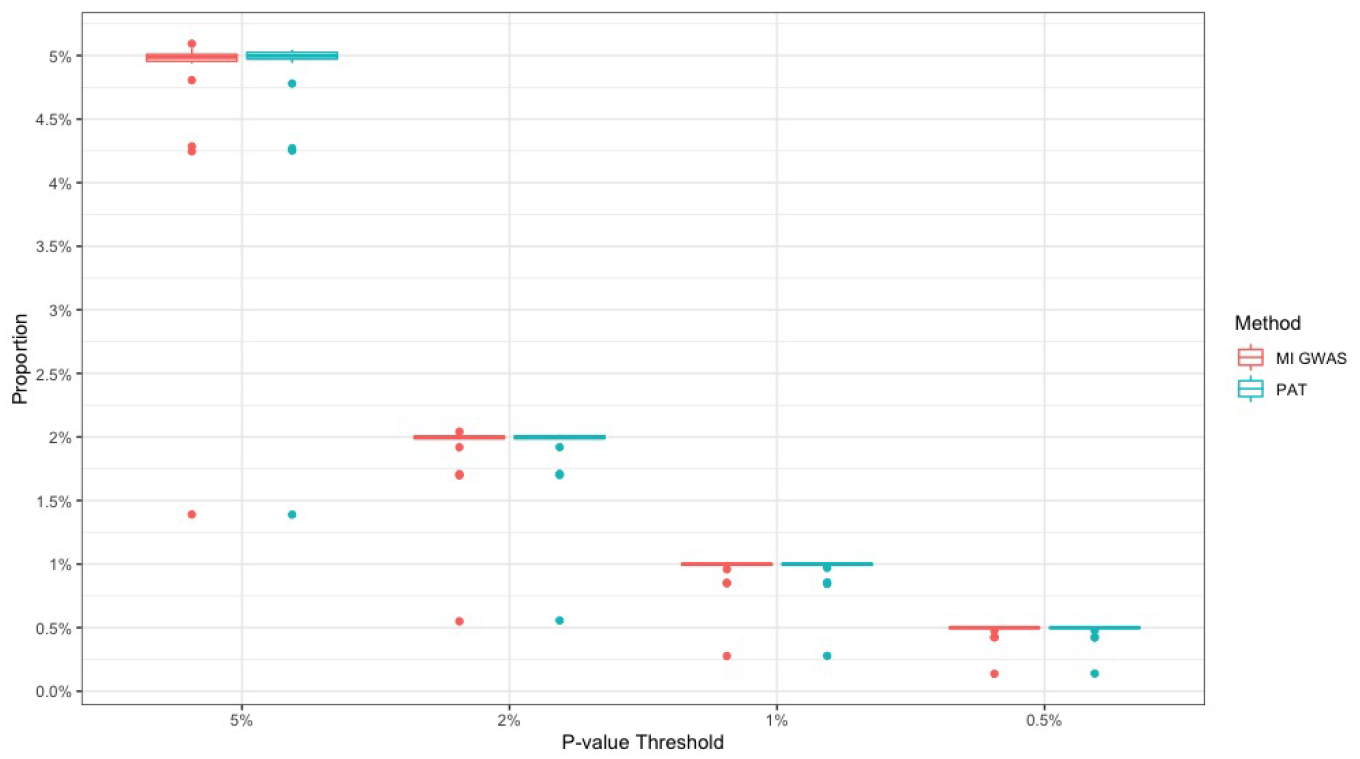
Comparison of false positive rate for PAT and MI GWAS. We use the UK Biobank data to estimate the genetic and environmental covariance matrices for four traits (body mass index, diastolic blood pressure, height, and systolic blood pressure) and inform PAT of the covariance structure between traits. We then simulate 10^8^ summary statistics 25 times with only environmental covariation between the traits. We examine the number of variants with p-values below 5%, 2%, 1%, and 0.5% level of significance to test how effectively the methods control false positives.

### 2.4 PAT increases power for pleiotropic effects

PAT is a likelihood ratio test whose rejection region is elliptical while the rejection region for MI GWAS is a square as shown in Fig. 2. Here, instead of comparing the shape, we use simulations to understand the relative power of each method when analyzing simulations based on four UK Biobank traits (see Appendix) [Neale Lab, 2017]. While PAT is able to account for overlapping sample sizes (see Methods), in these simulations individuals are assumed to be uniquely measured for each trait. This means there is no environmental correlation either in the simulations or when setting the test statistics for PAT (and MI GWAS). We make this assumption due to MI GWAS being unable to account for overlapping samples and to enable a fairer comparison of the methods.

To compare the power of MI GWAS and PAT, we simulate 10^8^ z-scores as if there is a genetic effect in every trait. The simulated genetic effect sizes begin by following the polygenic model and then for each subsequent set of 10^8^ z-scores the genetic effect sizes increases by a magnitude of 25. This is repeated 200 times until it is 5,000 times larger than the polygenic model. In Fig. 4, the power of MI GWAS and PAT are shown relative to the power of MI GWAS. When variants genetically affect all traits, PAT has more power than MI GWAS regardless of the genetic effect size. Furthermore, while power increases for both methods as effect sizes increase, PAT has a faster increase in power relative to MI GWAS. These results indicate that when pleiotropy is present, PAT has more power than MI GWAS to detect variants with weaker effect sizes.

**Fig. 4:**
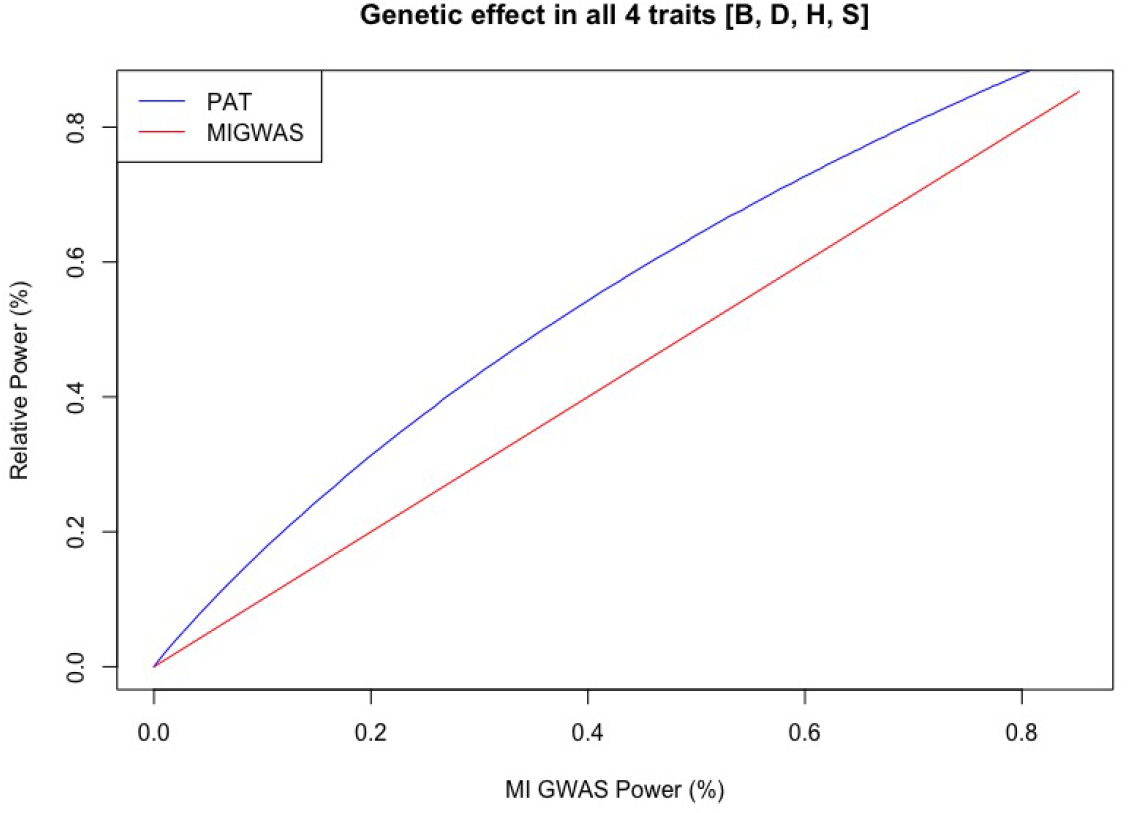
Comparison of relative power between PAT and MI GWAS when there is a genetic effect in every trait. We use the UK Biobank to estimate the genetic and environmental covariance matrices of four traits. The simulated effect sizes begin by following the polygenic model and then increase by a magnitude of 25 until the effect size is 5,000 times larger than the polygenic model. The power of PAT and MI GWAS to identify associated variants are shown relative to the power of MI GWAS.

### 2.5 PAT outperforms MI GWAS for many misspecified models of genetic effect

While PAT has more statistical power when the true underlying distribution of genetic effects is known, it is unreasonable to assume every associated variant affects all traits. We, therefore, test the robustness of PAT to model misspecificiaton. PAT is informed of two models, the null model (no genetic effect) and the full model (genetic effect in all traits). The simulations are based on four UK Biobank traits (see Appendix) but assume no environmental correlation. The z-scores are then simulated to violate the assumed alternative model by having a genetic effect in only a subset of the traits. The results can be seen in Fig. 5 where the relative power of PAT is compared to MI GWAS under various effect sizes.

**Fig. 5:**
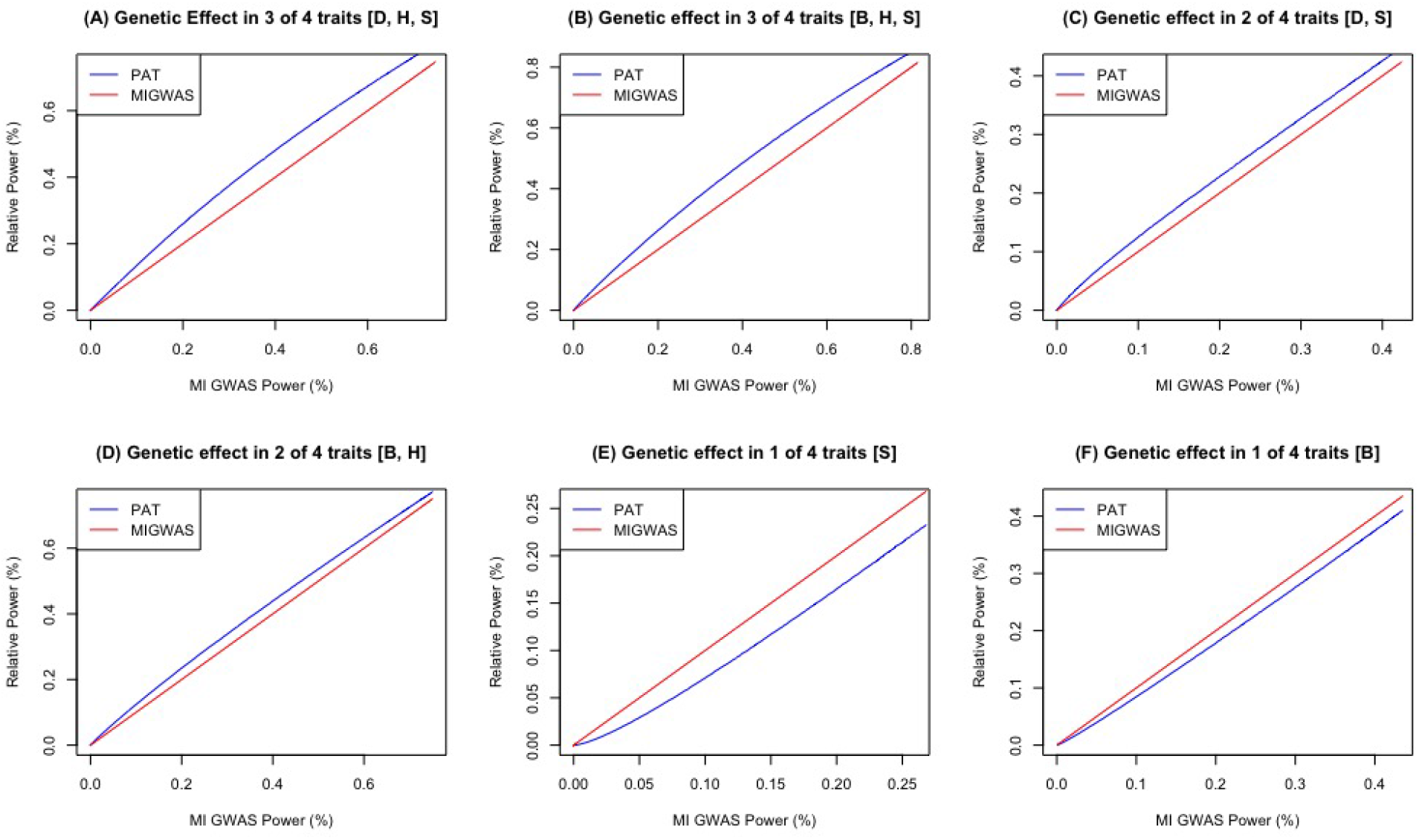
Comparison of statistical power when only a subset of traits have a genetic effect. These simulations are based off for four UK Biobank traits: body mass index (B), diastolic blood pressure (D), height (H), and systolic blood pressure (S). Each sub-figure show different sets of traits having their genetic effect set to zero. The relative performance of PAT is compared to MI GWAS with the power of MI GWAS as the baseline. In sub-figures A and B, the simulations model a genetic effect in three of the four traits while in sub-figures C and D, a genetic effect occurs in only half of the traits. In the last two sub-figures (E and F), there is a genetic effect in only one of the traits.

Fig. 5 explores the relative power for PAT and MI GWAS for six different configurations of effect. Sub-figures A and B model the case when there is a genetic effect in three of the four traits. For sub-figure A, the genetic effect size for body mass index is set to zero and for sub-figure B there is no genetic effect in diastolic blood pressure. Under both of these conditions, PAT has a substantial improvement in power over MI GWAS. For sub-figures C and D the simulations model a genetic effect in only two of the traits. In sub-figure C, the genetic effect size for body mass index and height are set to zero while in sub-figured D, there is no genetic effect for diastolic and systolic blood pressure. PAT is still more powerful than MI GWAS though the advantage is more modest. In the remaining two sub-figures (E and F) there is a genetic effect in only one trait. In sub-figure E, there is a genetic effect in systolic blood pressure while sub-figure F models there being an effect in only body mass index. When there is no pleiotropy, PAT is less powerful than MI GWAS. These simulations indicate that when pleiotropy is present, it is advantageous to jointly model genetic effects across traits, but this advantage decreases as the amount of pleiotropy decreases.

### 2.6 PAT is a power method for omnibus association testing in multi-trait GWAS

PAT has been shown to be a powerful method for detecting pleiotropic effects and is robust to many model misspecifications relative to a generalized version of single trait GWAS (MI GWAS); however, it is important to establish its performance relative to other multi-trait methods. Here, we compare three methods: PAT, HIPO, and MTAG [Qi and Chatterjee, 2018, Turley et al., 2018]. HIPO is an omnibus method that performs eigenvalue decomposition resulting in orthogonal components each of which is used to perform a weighted sum of z-scores. For this comparison, z-scores from all components are considered simultaneously and a variant is deemed associated as long as it is genome-wide significant for at least one component. The other method is MTAG, and it also uses a weighted sum of z-scores. MTAG, however, is not an omnibus method and instead computes and tests a z-score for each trait separately while leveraging information from the other traits. The results are converted to an omnibus test by determining if the variant is genome-wide significant for at least one trait with MTAG. There is no additional correction for multiple testing for MTAG or HIPO, and all methods are tested at *α* = 5 × 10^−8^.

In Table 1, 700,000 z-scores are simulated for four traits with the environmental and genetic covariance structure based on traits from the UK Biobank (See Appendix) [Neale Lab, 2018]. 10% (70,000) of the variants are causal in at least one trait. The first row in Table 1 corresponds to the 630,000 variants simulated under the null. All three methods correctly identify zero associated variants. The remaining 70,000 truly associated variants are equally split across seven configurations of genetic effect. For each of the configurations, there are three causal effect sizes: 35k, 21k, and 14k. The 10,000 variants for each configuration are split such that 5k, 3k, and 2k simulations came from each of the respective causal effect sizes. The seven configurations are subsets of the four traits, body mass index 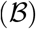, diastolic blood pressure 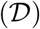, height 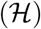, and systolic blood pressure 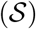.

**Table 1:** Comparison of multi-trait GWAS methods for omnibus testing. 700,000 variants were simulated with z-scores for four traits with 10% of variants being truly associated. The first column lists which trait has a genetic effect. The second column is the number of variants simulated under this specific model. The third column is the genetic effect size in the traits affected by the variant. The remaining three columns contain the number of variants identified as associated by an omnibus test by three method: PAT, HIPO, and MTAG.

| Genetic Effect | Number of Variants | $\frac{\sigma_g}{causal}$ | Genome-Wide Significant | | |
| --- | --- | --- | --- | --- | --- |
|  |  |  | PAT | HIPO | MTAG |
| No Trait | 630,000 | 0 | 0 | 0 | 0 |
| $\mathcal{B}, \mathcal{D}, \mathcal{H}, \mathcal{S}$ | 5,000 | 35,000 | **142** | 67 | 65 |
|  | 3,000 | 21,000 | **304** | 168 | 175 |
|  | 2,000 | 14,000 | **374** | 291 | 273 |
| $\mathcal{B}, \mathcal{D}, \mathcal{H}$ | 5,000 | 35,000 | **155** | 75 | 63 |
|  | 3,000 | 21,000 | **309** | 193 | 192 |
|  | 2,000 | 14,000 | **378** | 314 | 279 |
| $\mathcal{D}, \mathcal{H}, \mathcal{S}$ | 5,000 | 35,000 | **112** | 45 | 45 |
|  | 3,000 | 21,000 | **224** | 125 | 119 |
|  | 2,000 | 14,000 | **283** | 198 | 192 |
| $\mathcal{B}, \mathcal{H}$ | 5,000 | 35,000 | **144** | 72 | 70 |
|  | 3,000 | 21,000 | **228** | 139 | 139 |
|  | 2,000 | 14,000 | **338** | 257 | 246 |
| $\mathcal{D}, \mathcal{S}$ | 5,000 | 35,000 | 0 | 3 | **4** |
|  | 3,000 | 21,000 | 0 | **19** | 7 |
|  | 2,000 | 14,000 | 0 | **40** | 24 |
| $\mathcal{B}$ | 5,000 | 35,000 | 0 | **12** | 9 |
|  | 3,000 | 21,000 | 4 | **44** | 36 |
|  | 2,000 | 14,000 | 16 | **84** | 77 |
| $\mathcal{H}$ | 5,000 | 35,000 | **110** | 52 | 53 |
|  | 3,000 | 21,000 | **203** | 123 | 132 |
|  | 2,000 | 14,000 | **245** | 175 | 177 |
| Total | 700,000 | X | **3,569** | 2,496 | 2,377 |

While no simulation framework truly reflects the real world, this arrangement attempts to non-exhaustively model different scenarios that occur when analyzing z-scores from multiple traits. Namely, we explore the power to discover summary statistics of differing causal effect sizes and violations of a pleiotropic effect in all traits. Under these various configurations, none of the methods are powerful; however, PAT is the most powerful method in a majority of the scenarios simulated. This is especially true when there is a pleiotropic effect or when there is a genetic effect in a more heritable trait (e.g. height). In fact, across all scenarios PAT identifies 3,569 associated variants which is 43.0% increase over than the next best method HIPO (2,496). Across simulations, MTAG perform similarly to HIPO with a total of 2,377 variants identified. While PAT generally performs the best, both MTAG and HIPO do significantly better under some scenarios. One is when there is a genetic effect in traits that are both genetically and environmentally correlated such as diastolic and systolic blood pressure 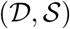. This condition is reflective of the simulations shown in Figure 2 particularly sub-figure (K). In this case, PAT is known to be conservative in the direction of environmental correlation. Lastly, HIPO and MTAG are both more powerful than PAT when there is no pleiotropy and the variant only affects body mass index 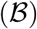. Overall, when pleiotropy is present, PAT is the most powerful method for omnibus association testing, but can be lose power in the presence of strong environmental correlation. These results are further explored on a per trait level (see below).

### 2.7 M-values provide accurate interpretation of omnibus association tests

As an omnibus test, PAT provide a single p-value for each variant tested across the set of traits. For the variants identified as genome-wide significant, we propose an extension to the m-values framework that enables a per trait interpretation (see Methods). Here, we use simulated data reflective of four real UK Biobank traits (see Appendix) to highlight the accuracy of m-values. In Fig. 6 (A), 1 million variants were simulated assuming there is a genetic effect in all four traits: body mass index, diastolic blood pressure, height, and systolic blood pressure. The second set of simulations shown in Fig. 6 (B) model a genetic effect in only body mass index and height. In Fig. 6, the truly associated traits are denoted with an asterisks (*) around the trait name. While the association test assumes the polygenic model, the data is simulated with the effect sizes equal to scaling *Σ_g_* and 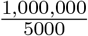. This results in the power being 50% in the first model and 44% when there is a genetic effect in only body mass index and height.

**Fig. 6:**
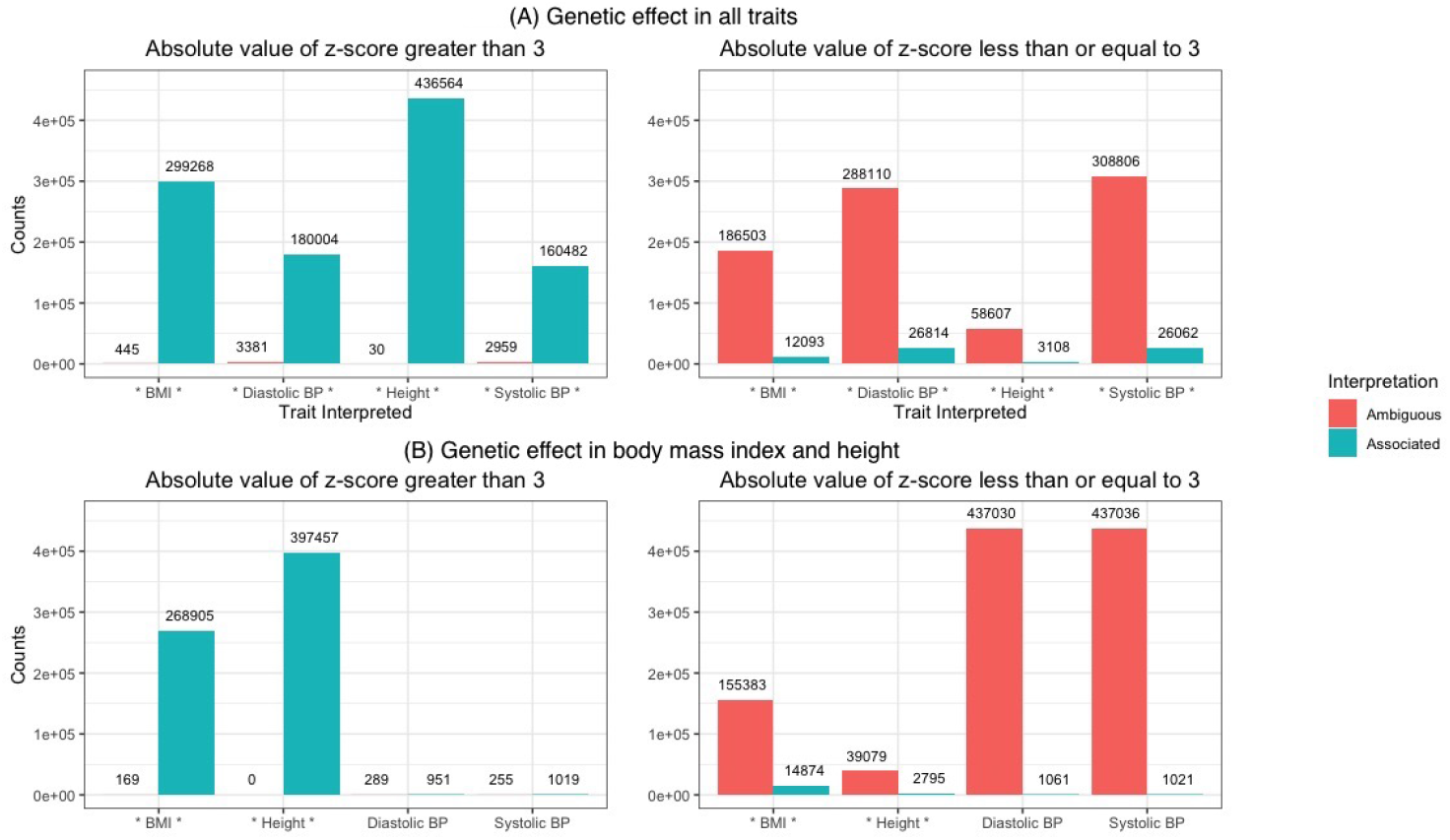
Interpreting per trait associations from omnibus significant variants. We simulate 1 million variants for four traits under two models. The first set of simulations assume there is a genetic effect in every trait (A), while the second model only has a genetic effect in body mass index and height (B). The associated traits are noted with an asterisks (*). The results for each trait are split based on the absolute value of the z-score and show the interpretation as either ambiguous or associated. The threshold for associated is an m-value greater than 0.9.

For every variant identified as associated, m-values are used to determine the variant has a genetic effect in each trait. This is done by enumerating over different configurations of genetic effect *Σ_g_*(*c*) where the configuration *c* = (1,1,1,1) is when there is a genetic effect in all traits (i.e. *Σ_g_*). When performing association testing, the variants are assumed to follow the polygenic model, however this model makes distinguishing between different configurations of genetic effect *Σ_g_*(*c*) for m-values difficult. As a result, we scale *Σ_g_*(*c*) to reflect the maximum likelihood estimate of the genetic effect size using the variants identified as associated (see Methods).

In Fig 6, the summary statistics (z-scores) for each trait are split according to whether their absolute value is greater than 3. This separation is done to distinguish cases where there is at least nominal signal in the trait from the instances where there may not be signal on the per trait level. When there is a genetic effect in all four traits and the summary statistic is greater than 3 (the first row left side), the m-value is greater than 0.9 for the vast majority of summary statistics. This means the variant was correctly interpreted as associated. Diastolic and systolic blood pressure had the most ambiguous associated variants with 3,381 and 2,959, respectively. This, however, is still less than 2% of the variants with at least a modest effect size (z-score greater than 3) being interpreted as ambiguous for each trait. Furthermore, when there is a modest effect size the overall false negative rate is 0.6% across the traits. When the absolute value of the summary statistic is less than or equal to 3, the interpretation is mostly ambiguous. This intuitively make sense because the z-score for many of the trait may not even be significant at the nominal threshold *α* = 0.05. Overall for diastolic and systolic blood pressure, more z-scores are less than or equal to three than are greater than three; however, even when the z-scores are small for diastolic and systolic blood pressure, more than 26,000 variants were correctly interpreted for both traits.

The second set of simulations models a genetic effect in only two of the traits (body mass index and height). When the z-score is greater than three, the m-value framework almost always correctly interprets when there is a genetic effect in body mass index and height. For body mass index only 169 of the variants were missed and none were left ambiguous for height. For diastolic blood pressure and systolic blood pressure, approximately 1,200 variants per trait have summary statistics with an absolute value greater than three. The m-value wrongly identified 951 and 1,019 of those variants as associated for diastolic blood pressure and systolic blood pressure, respectively. The same phenomenon happens when the z-score is less than or equal to three, and approximately the same number of variants for diastolic blood pressure and systolic blood pressure are misclassified as associated. Overall, 99.5% of the variants analyzed for diastolic blood pressure and systolic blood pressure are left with an ambiguous interpretation. For body mass index and height, 64.5% and 90.5% of all variants, respectively were correctly interpreted as associated when there is only a genetic effect in these two traits.

These simulations show the m-value framework has a low false positive assignment rate, and enables the correct classification of many associated variants. This is especially true when the z-score is greater than three. When the z-score is less than or equal to three, the interpretation is generally ambiguous regardless of if the variant is truly causal for the trait. While this is still a false negative assignment, many of these associations would fail to pass a nominal test for significance.

### M-value enable more per trait interpretations in multi-trait GWAS

PAT, MTAG and HIPO were previously compared in regards to their power to perform omnibus association testing (see above). Here, we investigate the per trait interpretation of these associations. As MTAG computes a p-value for every trait, the method provides a direct per trait interpretation. HIPO and PAT, on the other hand, perform an omnibus test; therefore, we will apply the m-value framework to assign a per trait association to the significant variants. For both methods, this is done by taking the associated variants and calculating the posterior predictive probability (m-value) of whether there is a genetic effect in each particular trait (see Methods). If the m-value is greater than 0.9, the variant is deemed associated with the trait. Otherwise, the interpretation is left ambiguous.

In Table 2, the comparison of PAT, MTAG and HIPO uses 700,000 simulations with 10% (70,000) causal variants equally divided into seven configurations of genetic effect. Under these seven configurations, the traits genetically affected by the variant vary. For each of the configurations, different effect sizes are also considered (see above). In Table 2, the number of per trait associations is reported for each trait under each model condition. When the variant does not truly have a genetic effect on the trait, the box is greyed to indicate false positives. Overall, Table 2 resembles the results shown in Table 1 which is to say when there is a pleiotropic effect PAT returns more associations. PAT is under powered relative to HIPO and MTAG, when strong environmentally correlation between traits is present or there is only an effect in one trait.

**Table 2:**
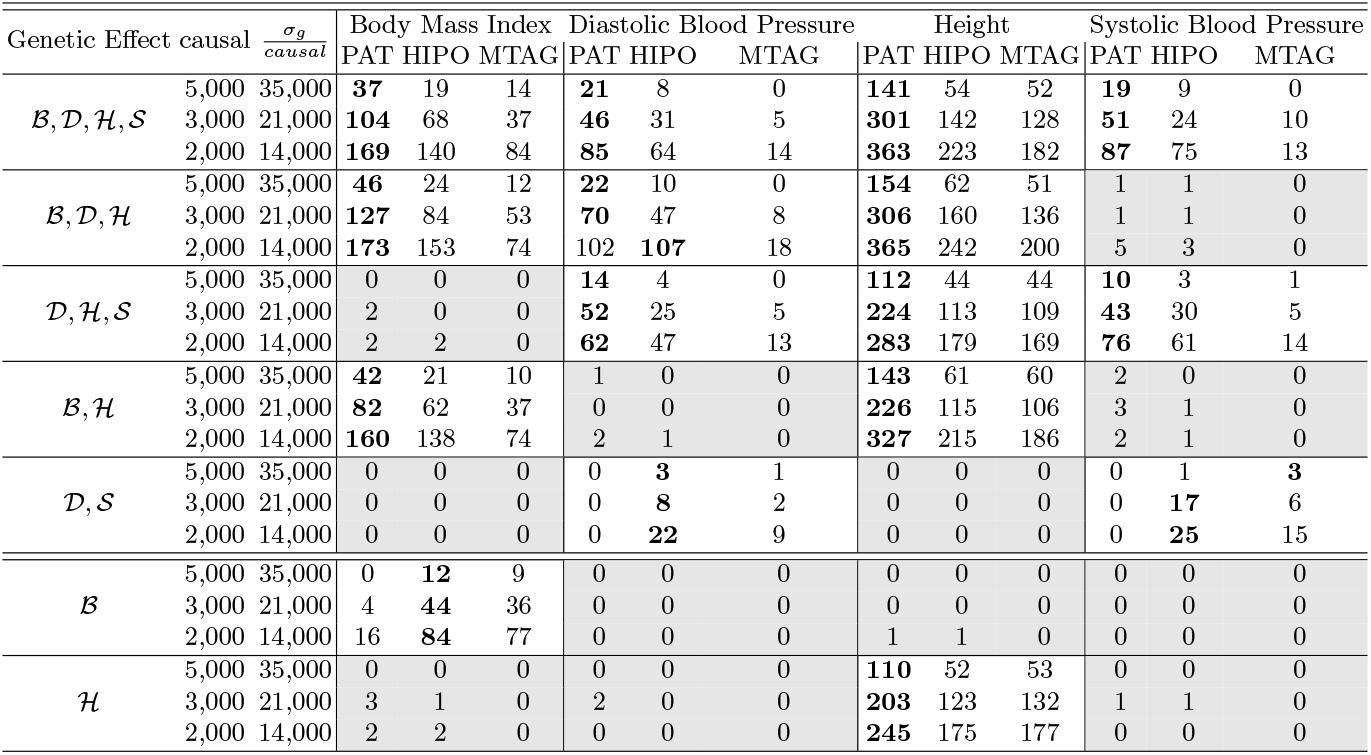
Comparison of multi-trait GWAS methods with per trait interpretation. 700,000 variants were simulated with z-scores for four traits with 10% of variants being truly associated. The first column lists which trait has a genetic effect. The second column is the number of variants simulated under this specific model. The third column is the genetic effect size of the variant. The remaining columns are split by trait where the performance of each method (PAT, HIPO, and MTAG) is shown per trait. These 12 columns show the number of variants identified as associated by each method for the specific trait. MTAG uses p-values while PAT and HIPO use the m-value framework to provide per trait associations.

While HIPO itself does not provide a per trait interpretation, applying m-values to its omnibus results consistently produces more per trait associations than MTAG. In fact, when there is a genetic effect in body mass index, diastolic blood pressure, and height and 2,000 causal variants 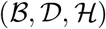, HIPO has more per trait associations than PAT for diastolic blood pressure. This means that while HIPO has less power than PAT for the omnibus test (314 and 378, respectively), it was able to provide the most per trait interpretations for this trait. This is due to HIPO identifying different associated variants than PAT which were then interpreted on a per trait level.

Overall, PAT identifies 5,223 true per trait associations from its 3,569 omnibus associations. For HIPO, the m-value framework interprets 3,430 per trait associations from its 2,496 associated variants. For PAT, this is 52.2% more true per trait associations than HIPO due to PAT having more power as an omnibus method. Finally, we consider MTAG which directly identifies 2,444 total per trait associations (2,377 omnibus associations). While HIPO and MTAG had similar power for the omnibus association, the m-value framework resulted in 40.3% increase in per trait interpretations. When comparing PAT and MTAG on a per trait level, PAT more than doubles the number of associations.

In Table 2, MTAG has no false positive per trait associations. The m-values produced for PAT and HIPO, however, do result in false positive assignments. PAT has 30 false per trait associations while HIPO has 15. This is 0.57% and 0.44% of their respective per trait interpretations. Even so, m-values are shown to have a low false positive assignment while enabling a significant increase in the true per trait interpretations. While powerful, it is key to acknowledge the m-value framework does not directly control false positives like a p-value threshold. This is because m-values are intended to provide empirical insights and interpretation to p-values not replace them. As a result, the current comparison between MTAG and the m-value interpretations is not an apples to apples comparison. In Supplementary Fig. 9, a ranked comparison between MTAG and m-values show that for any false positive rate, PAT and HIPO have more true positive per trait assignments than MTAG.

### PAT discovers novel per trait associations in the UK Biobank

While simulations have indicated PAT is a powerful method for association testing and m-values enable a per trait interpretation, we now apply this two step approach to real data to analyze the UK Biobank summary statistic for body mass index, diastolic blood pressure, height, and systolic blood pressure [Neale Lab, 2018]. Here, five methods are compared: Single Trait GWAS (how the z-scores and p-values were derived), MTAG, MI GWAS, HIPO and PAT [Turley et al., 2018, Qi and Chatterjee, 2018]. The set of variants are processed such that only variants which are biallelic, have non-ambiguous strands, a minor allele frequency greater than 1%, and an INFO score greater than 80% are retained. This leaves 7,025,734 variants that meet the criteria for all four traits. The reference and alternate allele are coordinated across traits by flipping the direction of the effect when necessary. LD-Score regression and cross-trait LD-Score regression are used to calculate the genetic and environmental covariance structure (see Appendix) [Bulik-Sullivan et al., 2015a,b]

Using the standard single trait GWAS, there were 211,546 uniquely associated variants across the four traits of interest. With MTAG, 164,263 uniquely associated variants were identified, 931 of which were novel associations. MI GWAS implicated 183,669 variants as associated, but none of the variants were novel discoveries due to MI GWAS having less power than single trait GWAS by design. When analyzing the traits with HIPO, 177,519 associated variants were found with 19,829 being new variants. PAT identified 200,112 uniquely associated variants with 22,095 being novel. None of the multi-trait methods identified more distinct variants than the standard single trait GWAS though MTAG, HIPO, and PAT identified new variants. This is likely due to insufficient power to capture variants associated with only one trait. For further exploration of the omnibus results see the Appendix.

When comparing the methods on their per trait associations, more associations are identified by leveraging multiple traits. While standard single trait GWAS identified 211,546 uniquely associated variants, only 18,764 were implicated as associated with more than one trait for a total of 233,540 associations as reported in Table 3. When analyzing the traits using MTAG, 18,054 out of 164,263 uniquely associated variants were found to be associated with more than one trait. This results in there being a total of 184,946 per trait associations. While single trait GWAS and MTAG provide a per trait p-value, MI GWAS, HIPO, and PAT do not. In order to interpret their associations, a per trait m-value must be assigned. When using the m-values framework, MI GWAS interpreted 325,113 per trait associations due to 96,519 of its 183,669 associated variants being associated to more than one trait. Out of the set of 183,669 uniquely associated variants, there were 8,213 whose interpretation was left ambiguous. This means that while those variants are significantly associated with the set of traits according to the omnibus test, the interpretation as to which of the traits it is specifically associated with is still ambiguous. HIPO identified 177,519 associated variants where 94,5333 were interpreted as associated with more than one trait. There were 862 with an ambiguous interpretation while 311,509 were interpreted as associated with at least one trait. Finally, the m-value framework is applied to PAT resulting in 363,955 per trait associations from the set of 200,112 unique variants. Out of which 111,126 variants were interpreted as associated with more than one trait and 9,869 were left with an ambiguous interpretation.

**Table 3:** UK Biobank data interpretation. We analyze four traits from the UK Biobank using five methods: Single Trait GWAS, MTAG, MI GWAS, HIPO, and PAT and show the variants associated with each trait. For Single Trait GWAS and MTAG, the per trait association is directly computed. For MI GWAS, HIPO and PAT, an omnibus association is first performed. The significant variants are then interpreted using the m-value framework, and m-values greater than 0.9 are interpreted as associated.

| Trait<br>Interpreted | Directly<br>From GWAS |  | M-value<br>>0.90 |  |  |
| --- | --- | --- | --- | --- | --- |
|  | Single Trait<br>GWAS | MTAG | MI GWAS | HIPO | PAT |
| body mass index ($\mathcal{B}$) | 37,205 | 32,527 | 64,706 | 65,462 | **67,139** |
| diastolic blood pressure ($\mathcal{D}$) | 18,593 | 17,610 | 56,369 | **58,294** | 56,271 |
| height ($\mathcal{H}$) | 160,227 | 117,882 | 155,730 | 136,519 | **191,420** |
| systolic blood pressure ($\mathcal{S}$) | 17,515 | 16,927 | 48,308 | **51,234** | 49,125 |
| Total | 233,540 | 184,946 | 325,113 | 311,509 | **363,955** |

We note that while MI GWAS cannot by definition have more power than Single Trait GWAS, once a variant is implicated as associated with at least one trait the interpretation could assign the association to multiple traits. This means that as long as the effect size in one variant is large enough to result in MI GWAS finding the variant significant, the weaker effect sizes can still be interpreted using m-values. This is because the m-values framework leverages the genetic and environmental covariation between traits regardless of whether or not the original method models it. As a result, the m-value framework enables an increase in per trait associations. In fact, PAT has over 100,000 more per trait associations than single trait GWAS in Table 3 even though it implicated fewer variants. For body mass index, PAT, MI GWAS and HIPO almost double the number of per trait associations and nearly triple it for systolic blood pressure. For diastolic blood pressure, the number of per trait associations is more than tripled due to the m-value framework. In Table 3, MTAG performs on par with single trait GWAS on a per trait level. One reason for the difference in performance is the nature of the methods. For MI GWAS, HIPO, and PAT, the variant is first implicated and then interpreted on a per trait basis while MTAG and single trait GWAS assign statistical significance for each trait separately.

### Analyzing PAT’s novel associations in the GIANT consortium

PAT identified 22,095 novel associations when simultaneously analyzing four traits from the UK Biobank. All associated variants had a per trait interpretation to at least one trait. The breakdown of the per trait associations is as follows: 12,261 variants were interpreted as associated with body mass index, 7,868 with diastolic blood pressure, 21,119 with height, and 7,605 were interpreted as associated with systolic blood pressure.

Now equipped with novel per trait associations, these discoveries should be validated by an external dataset. We use summary data from the GIANT consortium to see if any of the new associations for body mass index and height can be reproduced [Wood et al., 2014, Locke et al., 2015]. For body mass index, the European summary statistics from the GIANT consortium contains 2,554,638 SNPs. Out of the 12,261 novel variants interpreted to be associated with body mass index in the UK Biobank, 3,946 were found in the GIANT consortium by matching on the RSID, reference, and alternate allele and have a minor allele frequency reported. In the cases that the reference and alternate allele differs between the two data sets, the direction of the effect size in the replication data set is flipped. After identifying which variants are present in both data sets, the set of variants is pruned by taking the largest m-value (i.e., posterior predictive probability) and removing all other variants within a 1MB region. We prune on the m-value instead of the p-value due to the significance of PAT’s p-value potentially being driven by a different trait. This clumping algorithm results in 408 variants being tested for replication in body mass index. This process is repeated in height where the GIANT consortium has 2,550,859 variants to be considered. Out of the 21,119 variants that were interpreted as associated with height in the UK Biobank, there are 7,068 which can found in the GIANT data set. After clumping these variants to the peak m-value per megabase region, there are 735 independent associations.

In order to test the replication rate, we perform a one-sided z-test in the direction of the effect size (β) in the UK Biobank. For body mass index, 378 out of 408 variants (92.6%) have the observed effect sizes between the UK Biobank and GIANT consortium in the same direction. We test each variant for replication using the level of significance 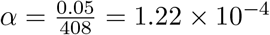 where 14 of the tested variants have a p-value below the threshold for replication. For height, 97.4% (716) of the tested variants have the effect sizes in the same direction between the UK Biobank and the GIANT consortium. For replication, we set the level of significance to 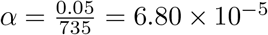. Here, we see that 197 of the 735 variants replicate.

For the variants that failed to replicate, there are a number of reasons this may have occurred. In Table 4, we explore how statistical power affects our replication rate. We first bin the variants into deciles by their replication power. For each decile, we calculate the average power this set of variants has to replicate the effect size observed in the UK Biobank in the GIANT consortium. We note that the GIANT consortium does not release the minor allele frequency observed in their samples but instead provides the minor allele frequency observed in HapMap [Frazer et al.]. While a reasonable estimate inaccuracies in the minor allele frequency may impact the estimate of power. The average power over all variants for body mass index is 39.4%, however we only see a replication rate of 3.4%. For height, the replication rate is 26.8%, but the overall power is 65.5%. In table 4, we see for both traits that as power increases so does the replication rate; however, neither trait replicates at the expected rate. The only exception is that for height when the power is between 0-30%, we see a replication slightly over the expected rate.

**Table 4:** Replication power in the GIANT consortium. Using novel associations found in the UK Biobank by PAT for body mass index and height, associations are clumped using the lead variant as determined by the m-value. For each variant, we calculate replication power and bin the variants into deciles. The first column lists the trait. The second column is the decile while the third column is the average power within the set. The number of variants tested for replication, the expected number of replications, and the number of variants that replicated are reported in the next three columns, respectively. The final column contains the number of variants with effect sizes from the GIANT consortium in the same direction seen in the UK Biobank. A binomial test on whether the proportion of effect sizes in the same direction across studies is greater than 50% of all tested variants in the set. A single asterisks in the final column means the results are significant at the nominal *α* = 0.05 and two asterisks indicates significance at 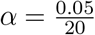.

| Trait | Power | Average Power | Number of Variants Tested | Expected Number of Replications | Number of Replications | Number of Effect Sizes in Same Direction |
| --- | --- | --- | --- | --- | --- | --- |
| Body Mass Index | 0-10% | 6.8% | 44 | 3 | 0 | 35** |
|  | 10-20% | 15.2% | 92 | 14 | 0 | 84** |
|  | 20-30% | 24.7% | 57 | 14 | 1 | 54** |
|  | 30-40% | 35.5% | 40 | 14 | 0 | 37** |
|  | 40-50% | 45.3% | 33 | 15 | 2 | 32** |
|  | 50-60% | 55.5% | 23 | 13 | 0 | 23** |
|  | 60-70% | 64.5% | 43 | 28 | 5 | 40** |
|  | 70-80% | 75.2% | 53 | 40 | 1 | 51** |
|  | 80-90% | 85.9% | 15 | 13 | 1 | 15** |
|  | 90-100% | 92.6% | 8 | 7 | 4 | 7* |
| Height | 0-10% | 4.2% | 15 | 1 | 1 | 12* |
|  | 10-20% | 13.4% | 11 | 1 | 2 | 11** |
|  | 20-30% | 24.7% | 13 | 3 | 4 | 12** |
|  | 30-40% | 36.0% | 30 | 11 | 5 | 26** |
|  | 40-50% | 45.7% | 46 | 21 | 5 | 46** |
|  | 50-60% | 55.5% | 90 | 50 | 27 | 89** |
|  | 60-70% | 65.8% | 135 | 89 | 38 | 130** |
|  | 70-80% | 75.0% | 303 | 227 | 92 | 300** |
|  | 80-90% | 83.1% | 75 | 62 | 21 | 73** |
|  | 90-100% | 93.0% | 17 | 16 | 2 | 17** |

While we have shown that our replication rate is below expectation, the expected replication rate is likely overestimated. This is due to an inflation in the discovery cohorts effect size; a phenomenon known as winner’s curse [Palmer and Pe’er, 2017]. As the variants we are testing for replication were not identified as associated by the original single trait GWAS, these variants have sub-optimal power for discovery. This means these variants have small effect sizes and were only found associated after leveraging their covariance structure with other traits. As a result, the bias in the effect size will be much larger here than in variants that were already well powered for discovery. For the non-replicating variants, further power increases are essential to better tease out which variants warrant follow up analyses.

While many variants failed to replicate potentially due to insufficient power or winner’s curse, we also tested whether the effect sizes were in the same direction between GIANT and the UK Biobank. This is tested because if the effect sizes were truly zero, the concordance of effect size across the data sets is expected to be 50%. If the effect size is truly non-zero, a higher concordance across data sets is expected. We test each decile of power using a binomial test to determine whether the proportion of effect sizes in the same direction is greater than 50% of the variants. Using the significance threshold *α* = 0.05, every test was significant. As there were 20 tests, we adjust for multiple testing using a Bonferroni correction 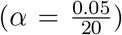. All but two tests were statistically significant at this new threshold. A test of overall concordance in body mass index tests whether 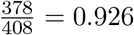 is statistically significantly greater than 0.5 returns a p-value of 4.34 × 10^−78^. We also test height for whether 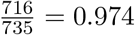 is greater than 0.5 which returns a p-value equal to 1.06 × 10^−184^. Therefore, we conclude that while the actual replication rate is low, there is still some evidence of there being real genetic signal in these variants.

### Computational speedup with importance sampling

We now show how the cost of null simulations can be reduced using importance sampling. When setting the critical value *κ* for PAT’s likelihood ratio test, the data is simulated according to the null distribution 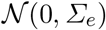. As a result, a likelihood ratio greater than *κ* is expected only *α* × *n* times. As GWAS uses the significance threshold of *α* = 5 × 10^−8^, the number n needs to extremely large to ensure replication of results; in practice n = 10^10^. Simulating and storing 10^10^ vectors of summary statistics is computationally expensive, especially in terms of memory. This burden can be reduce using importance sampling where means the null data is simulated according to a different distribution 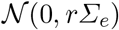 where *r* is a scaling factor that increases the number of samples that are significant. We note that importance sampling adjusts the weights of the samples in estimating the p-values (see Methods). If r is well chosen *κ* can be set with fewer simulations.

In Table 5, the critical value *κ* is estimated 25 times and the sample variance of these estimates provides a measure of the stability of the sampling. This is repeated for different values of the scaling factor r and number of samples *n*. We define the ratio of the sample variance using importance sampling to the sample variance of null simulations as stability. When the ratio is close to one, the estimated *κ* using importance sampling is as stable as the *κ* estimated directly using null simulations and values larger than one indicate importance sampling has a smaller variance. Four traits: body mass index 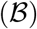, diastolic blood pressure 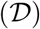, height 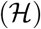, and systolic blood pressure 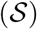 are considered in these simulations as well as subsets of the traits. When using 10^6^ simulations, we find using importance sampling is consistently more stable for all reported scaling factors, *r*. For diastolic blood pressure and systolic blood pressure, importance sampling is also more stable for all reported scaling factors using 10^5^ simulations. When using the scaling factor *r* = 8 for *n* = 10^5^ simulations, the variance for the value *κ* when using importance sampling is approximately equal to the variance using 10^10^ null simulations across the various sets of traits. For most sets of traits importance sampling is still slightly more stable; it is only for body mass index, height, and systolic blood pressure that it is less stable with a ratio of 0.93 which is still very close to 1. This means, the same stability can be achieved using only 10^5^ simulations which is 10^5^ fewer simulations. This reduction in computational resources holds true across the data sets. In practice, however, the use of 10^6^ simulations is more practical as the stability of the critical value *κ* is less sensitive to the setting of *r*. This still results in using 10,000 fewer simulations.

**Table 5:** Stable estimates of critical values in fewer null simulations. We generate the critical value *κ* at *α* = 5 × 10^−8^ 25 times for various combinations of four traits: body mass index 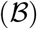, diastolic blood pressure 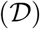, height 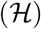, and systolic blood pressure 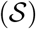. We simulate data according to 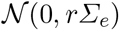 for *r* = {5, 6, 7, 8} and for *n* = 10^4^, 10^5^ and 10^6^ simulations. We then take a ratio of the variation in the estimated critical value *κ* which we call the stability. The first column is the set of traits and the variance for 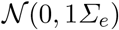 using *n* = 10^ιν^ simulations. The second column is the number of simulations while the remaining columns show the stability for different scaling factors of the covariance matrix *r* : *r* = {5, 6, 7, 8}.

| Set of Traits<br>(Variance) | Number of<br>Simulations | Scaling of Covariance Matrix ($r \times \Sigma_e$) | | | | | |
| --- | --- | --- | --- | --- | --- | --- | --- |
|  |  | 5 | 6 | 7 | 8 | 9 | 10 |
| $\mathcal{B}, \mathcal{D}, \mathcal{H}, \mathcal{S}$<br>(9e-07) | 1e6 | 6.94 | 13.82 | 6.88 | 12.10 | 6.49 | 6.12 |
|  | 1e5 | 0.741 | 1.01 | 1.40 | **1.01** | 0.58 | 0.70 |
|  | 1e4 | 0.046 | 0.12 | 0.13 | 0.06 | 0.09 | 0.10 |
| $\mathcal{B}, \mathcal{D}, \mathcal{H}$<br>(5e-07) | 1e6 | 5.27 | 4.92 | 7.50 | 7.11 | 13.76 | 7.22 |
|  | 1e5 | 0.54 | 0.53 | 0.58 | **1.15** | 1.00 | 0.69 |
|  | 1e4 | 0.035 | 0.045 | 0.045 | 0.074 | 0.096 | 0.08 |
| $\mathcal{B}, \mathcal{D}, \mathcal{S}$<br>(2e-07) | 1e6 | 11.11 | 7.93 | 9.35 | 8.32 | 11.10 | 12.07 |
|  | 1e5 | 0.62 | 0.81 | 1.14 | **1.17** | 0.90 | 0.94 |
|  | 1e4 | 0.10 | 0.08 | 0.16 | 0.13 | 0.17 | 0.18 |
| $\mathcal{B}, \mathcal{H}, \mathcal{S}$<br>(4e-07) | 1e6 | 3.20 | 11.75 | 5.53 | 6.22 | 9.29 | 6.08 |
|  | 1e5 | 0.33 | 0.65 | 0.64 | **0.93** | 0.63 | 0.73 |
|  | 1e4 | 0.07 | 0.06 | 0.09 | 0.07 | 0.09 | 0.22 |
| $\mathcal{D}, \mathcal{H}, \mathcal{S}$<br>(9e-07) | 1e6 | 7.23 | 7.97 | 18.51 | 18.31 | 16.39 | 12.73 |
|  | 1e5 | 0.54 | 0.73 | 0.92 | **1.30** | 1.26 | 1.47 |
|  | 1e4 | 0.10 | 0.08 | 0.11 | 0.08 | 0.09 | 0.31 |
| $\mathcal{B}, \mathcal{H}$<br>(6e-07) | 1e6 | 10.67 | 12.22 | 22.48 | 22.551 | 20.586 | 18.295 |
|  | 1e5 | 0.59 | 0.83 | 0.88 | **1.72** | 3.02 | 2.04 |
|  | 1e4 | 0.10 | 0.13 | 0.20 | 0.16 | 0.22 | 0.27 |
| $\mathcal{D}, \mathcal{S}$<br>(6e-08) | 1e6 | 28.83 | 21.53 | 110.27 | 27.46 | 33.39 | 72.58 |
|  | 1e5 | 1.492 | 4.04 | 3.49 | **4.58** | 7.39 | 5.05 |
|  | 1e4 | 0.18 | 0.37 | 0.40 | 0.63 | 0.51 | 0.67 |

## 3 Discussion

Here, we present PAT, a method that leverages pleiotropy for joint association testing in multiple traits as well as an extension to the m-value framework. Through simulations, PAT is shown to control the false positive rate as well as significantly increase statistical power to detect pleiotropic effects. The impact of misspecifying model parameters on PAT is also explored. In general, PAT is robust to there being a genetic effect in only a subset of traits. Additionally, when setting the critical value using null simulation, the computational burden is greatly reduced through importance sampling. One major limitation of PAT is its lack of per trait interpretation. This is overcome by the extension to the m-values framework presented here. It enables a per trait interpretation of PAT and other omnibus association tests. Through simulations, we find that the false positive assignment rate from m-values is low.

Additionally, PAT is compared to two current methods: MTAG and HIPO. In simulations, MTAG and HIPO are shown to have similar levels of statistical power while PAT is shown to be a more powerful method for omnibus association testing. By interpreting the associated variants using the m-value framework, PAT was also able to identify more per trait associations as well. Even so, there were some scenarios where PAT is underpowered such as when the variants are greatly effected by environmental correlation. In these cases, MTAG and HIPO lend a better model for joint analysis of traits. When traits of interest are all moderately to highly genetically and environmentally correlated such as the diastolic and systolic blood pressure, a method such as MTAG or HIPO may be preferred to PAT due to its conservative nature in the presence of strong environmental correlation.

In addition to simulations, PAT analyzes four real traits in the UK Biobank discovering 22,095 novel associations. After computing m-values and clumping the per trait associations, the replication of associated variants in body mass index and height are tested in the GIANT consortium. For body mass index, 14 of the 408 tested variant replicated. The replication rate in height was much higher with 197 out of 735 variants replicating. While the replication rate is below expectation, this may be due to winner’s curse. The variants testing for replication are novel variants that were under powered in the original association. This means these newly discovered associations have small effect sizes and will, therefore, be more affected by the bias in effect sizes [Palmer and Pe’er, 2017]. In addition to testing replication, we tested whether the effect sizes between the UK Biobank and GIANT are in the same direction. Overall, there is significant evidence that the effect sizes are in the same direction which is improbable if all of the effect sizes are truly zero. Further increases in power are needed to determine which of the non-replicating variants are providing true signals of genetic effect.

Through simulations and real data analysis, PAT is shown to be an effective method for leveraging pleiotropy for joint analysis of GWAS summary statistics. However, the optimal number of traits to jointly analyze is not explored. As the number of traits jointly analyzed increases, the genome-wide estimate of genetic correlation will cease to hold across all traits. This would result in fewer novel associations as the power gains would be stunted by model mispecification. Further exploration is needed to determine which traits should be analyzed together and how to effectively cluster the traits into these sets.

Another limitation to PAT is the assumption of the genetic covariance structure being constant across the genome. Modeling the local genetic covariance structure would increase power to identify associated variants. This would need to be done with care as to not over fit the alternative model in that region. If done well, however, this should lend even more power to PAT as it could better reflect the covariance structure between the z-scores [Shi et al., 2016, 2017].

## 4 Method

### 4.1 Association testing in a single quantitative trait (GWAS)

We now describe the standard approach for determining if a genetic variant g is associated with a quantitative trait y. Let *y* and g be measured for *N* individuals where *g_j_* ∈ {0,1,2} is the minor allele count for each individual *j*. The column vector g is then standardize according to the population proportion of the minor allele *p* where 2*p* is the mean and 2*p*(1 − *p*) is the variance of *g*. This standardized column vector *x* is defined as follows 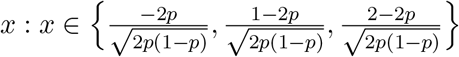. The quantitative trait y is normally distributed such that 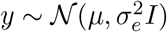 where *μ* is the mean and 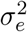 is the variance of the trait. This can then be mean-centered and scaled which results in the column vector 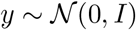. We can now assume the following linear model:

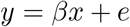

where *β* is the effect size of the variant *x* on the trait *y* and the residuals e follow the standard normal [Eskin, 2015]. Ordinary Least Squares results in the estimator 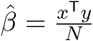 where 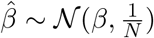. Setting 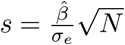 results in the following Gaussian: 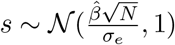.

We now test the null hypothesis: *x* is *not* associated with *y*. More formally this tests if *β* = 0 or 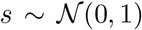. The null model is rejected if |*s*| > *z* where *z* is the z-statistic at the *α* level of significance for the standard normal distribution. The corresponding critical value 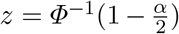. Typically, human GWAS uses *α* = 5 × 10^−8^ [Consortium, 2005, McCarthy et al., 2008, Pe’er et al., 2008].

### Generalizing GWAS testing to multiple traits (MI GWAS)

As previously stated, GWAS traditionally analyzes each trait *y_i_* in a set of T traits independently. In fact, each trait may be measured on distinct sets of individuals. Let us assume none of the traits *y*_1_,.., *y*_T_ have overlapping individuals; therefore every trait *y_i_* and the standardized genetic variant *x* is measured for *N_i_* individuals. This assumption will later be relaxed. For now, the z-score for trait i, s¿, is tested for whether |*s_i_*| > z, and this process is repeat for each trait independently. Another approach instead of instead of performing T different hypothesis tests is to determine whether the variant is associated with at least one of the traits. The corresponding null hypothesis is the variant is not associated with any of the traits. We refer to this method as multiple independent GWAS (MI GWAS). This results in *β*_1_ = … = *β_T_* = 0 which is equivalent to saying the null model is 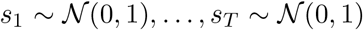. A simple way to test our null hypothesis is to check if the largest *s_i_* ∈ *S* = {|*s*_1_|,…, |*s_T_*|} is greater than the critical value z though z will now need to be corrected for multiple testing. This can be done using a Bonferroni correction for the number of traits, *T*, so the critical value is 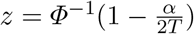.

Another method for setting the critical value is using null simulations. This is done by simulating data according to *S* = {*s*_1_,…, *s_T_*} such that every *s_i_* is under the null hypothesis. As all traits are measured for different groups of individuals, there is no covariation between any pairs of traits. This means the multivariate 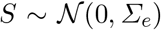 has the identity matrix as its covariance matrix; therefore, we simulate 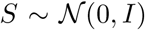 n times keeping the *max*{|*s*_1_|,…, |*s_T_*|} for each S. We then sort the n retained values and assign a p-value to each critical value using the quantile.

### 4.3 Using pleiotropy for association testing in multiple traits (PAT)

Another method for hypothesis testing is a likelihood ratio test which compares the null model to a proposed alternative model. Currently, only the null model has been defined. For a single quantitative trait y whose null hypothesis is *β* = 0 and 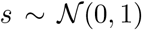, the alternative hypothesis is *β* ≠ 0 and *β* is assumed to follow a Gaussian distribution: 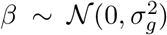, where 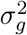 is the additive, per-SNP heritabilty of the trait. As Gaussian distributions are conjugate priors to Gaussian likelihood functions, the distribution of *β* can be used to get the Gaussian posterior predictive distribution, 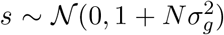. Two models that describe s have been defined and result in the following likelihood ratio:

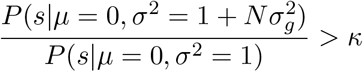

If the ratio of the likelihood functions is larger than *κ*, the null hypothesis is rejected. Before expounding on how to set *κ*, we will first extend the likelihood ratio test to the case of multiple traits.

We retain the assumption that the traits are not measured on the same individuals. This means, there is no environmental correlation, so under the null hypothesis 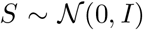. The assumption about distinct sets of individuals does not, however, have the same implication for the genetic correlation between genetic effects. Letting the cov(*β_i_,β_k_*) = *σ_gik_* we can derive 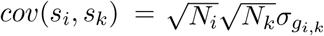. This results in the alternative model being:

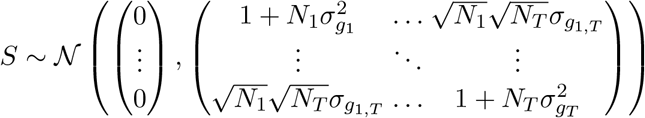

which can be written as 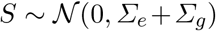. This means under the alternative model, the covariance of S is the sum of the environmental and genetic covariance where for now the environmental covariance is still the identity matrix, I. The likelihood ratio for PAT is now defined as:

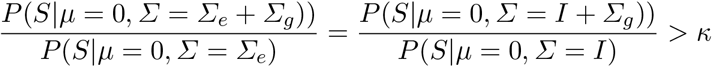

The critical value *κ* is set for PAT using the same null simulations of 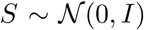. This time the likelihood ratio for each S is retained, sorted, and assigned a p-value using the quantile.

### 4.4 Overlapping Individuals for Multiple Traits

We now relax the assumption that no individual is measured for more than one trait. Under the null hypothesis, this means that *Σ_e_* in 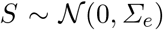 would not be the identity matrix I. In this case, 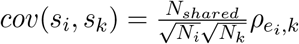. This means the covariance between *s_i_* and *s_k_* is the environmental correlation between the traits, and the environmental correlation is weighted by the proportion of overlapping individuals. Under the alternative hypothesis, we have 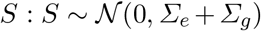. We note that while sample overlap between traits affects *Σ_e_*, it does not impact *Σ_g_*.

### 4.5 Importance Sampling for Null Simulations

When performing null simulations, the number of simulations n must be large enough that the critical value *κ* is stable across estimates. In practice, this can require n to be very large when *α* is really small because simulating 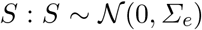 with a likelihood ratio larger than *κ* is expected to occur *α* × n times. One method for reducing the number of simulations is importance sampling.

To explain our approach we first review importance sampling in one trait. While, traditionally 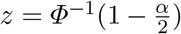 is used to set the critical value *z* for the standard normal. It is also possible to use null simulations just as we do for MI GWAS and PAT. We simulate 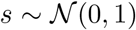 n times and sort |*s*| and assign the p-values using the quantile. To obtain the significance of a specific critical value such as 5.2, enough null simulations must be performed to have a sufficient number of samples above the critical value. The p-value would then be estimated by counting the number of samples above the critical value divided by the total number of samples. Unfortunately, for very significant p-values this requires a very large number of samples since the vast majority of samples are below the critical value.

Importance sampling reduces the number of simulations needed for setting the critical value by simulating data according to a different distribution *v* where *v* results in samples larger than the critical value *z* to occur more frequently. The procedure for estimating the p-value will then be adjusted to account for the differences between the two distributions, *s* and *v*. In our approach, *v* has the following distribution 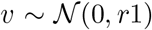, and in Fig. 7 the scaling factor r = 2 is used. In this figure, the critical value *z* ≈ 1.96 for *α* = .05 is shown for the null distribution s. We can see in Fig. 7 that the distribution *v* has many more samples in the tails; therefore, the significance level *α* does not correspond to *z* for the distribution *v*. The p-value using importance sampling is estimated for each data point by first computing a weight w. This weight *w* is the likelihood ratio 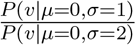 of the data points from *v* under the two models. By summing the weights of samples larger than the critical value and dividing by the sum of the weights for all samples, the p-value can be set for each critical value. We note that if *r* = 1, then *s* and *v* are identical and all the weights are 1. In this case, importance sampling and the standard approach are equivalent.

**Fig. 7:**
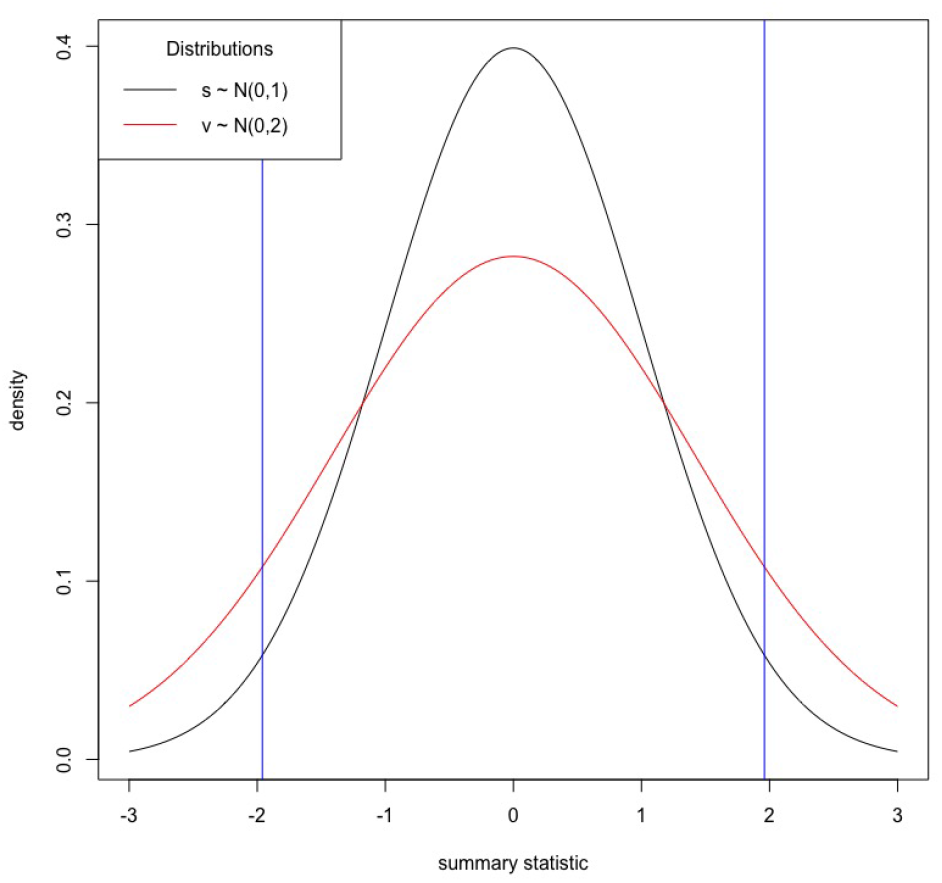
Using importance sampling for setting critical values. We simulated data according to two univariate Gaussian distributions 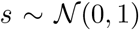 and 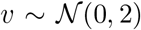 show the densities. We show the critical value *z* ≈ 1.96 for *α* = 0.05. We would expect to see the critical value |*z*| or larger more often when simulating data according to *v* than when simulated under the distribution of s.

We can now extend this to learning about the null distribution of 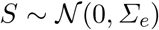 by simulating data according to the distribution 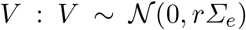. Again we will find that a well chosen alternative distribution *v* results in more statistics greater than *κ* in fewer simulations. The weight *w* is the likelihood ratio 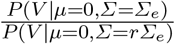 and will be used to obtain p-values as described above.

### 4.6 Interpreting GWAS meta-analyses

When performing an omnibus hypothesis test, there is only one p-value which cannot be directly interpreted on a per trait level. In previous work, the statistic m-values were introduced to enable interpretation of GWAS meta-analyses across studies, with m-values being the posterior probability of a genetic effect per study. [Han and Eskin, 2012]. The original m-values assumed that across studies the effect sizes are similar as it considered the same trait across multiple studies. When applying this framework to multiple traits, the model needs to account for differing effect sizes to prevent spurious results. Below, we describe the extension to the m-value framework which assumes a random effects model for the genetic effect and that the effect sizes reflect the genetic correlation estimated genome-wide.

We assume there are T traits for which a variant has been identified as associated by PAT (or another omnibus test). While the variant is known to be associated, there are many possible configurations of an effect. There may be an effect in all traits in which case the configuration is *c* = (1,…, 1), or there may only be an effect in the first trait, *c* = (1, 0,…, 0). The set of all configurations can be written as *c* = {0,1}^T^ where |*C*| = *2^T^*. For each trait i, there is subset of configurations *C_i_* ⊂ *C* that are compatible with the variant having a genetic effect in that particular trait, where |*C_i_*| = 2^T-1^. This means that every *c* ∈ *C_i_*, the ith index is always 1.

When PAT determines a variant is associated with the set of T traits, it assumes the variant affecting all T traits; therefore, the assumed posterior predictive distribution of effect is *P*(*S*|*μ* = 0, *Σ* = *Σ_g_* + *Σ_e_*). While pleiotropy is ubiquitous, the assumption that every variants affects all traits is not realistic. We will now define the genetic covariance matrix *Σ_g_*(c) that corresponds to a configuration *c* where

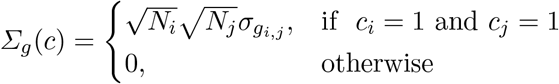

We note that when *c* = (1,…, 1), *Σ_g_* = *Σ_g_*(*c*). With this in mind, m-values works by summing the posterior probabilities that corresponding to the configurations in *C_i_* and dividing by the the total sum of all posterior probabilities (set of configuration in C). Therefore, for each trait *i*:

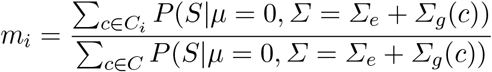

where *s* are summary statistics across the *T* traits for one variant, and the m-value *m_i_* is the proportion of the all posterior probabilities compatible with there being an effect in trait i. When *m_i_* > 0.9, the variant is assumed to be associated with the ith trait. Otherwise, the interpretation is left ambiguous.

While the assumed covariance structure of *Σ_g_* estimated genome-wide follows the polygenic model, for interpretation purposes this assumption is relaxed. Under the polygenic model, every variant has an effect; therefore, the expected effect size of each variant is 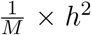 where *M* is the total number of variants and *h*^2^ is the estimated additive heritability of the trait. When only considering the variants found genome-wide significant, the expected effect size of these variants needs to be to estimated. We do this by estimating the number of causal variants *Q* and rescale the genetic covariance matrix *Σ_g_* by 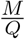 for the m-value interpretation framework.

This is necessary because *h*^2^ ∈ [0,1] and with genome-wide association studies using millions of variants, *Σ_g_ + Σ_e_* ≈ *Σ_e_* under the polygenic model. While a valid model for association testing, distinguishing between different configurations of *Σ_g_*(*c*) to calculate the m-value is very difficult. Therefore, we scale *Σ_g_* and the resulting *Σ_g_*(*c*) by randomly selecting one associated variant per 100KB region for a total of *k* variants. We then perform a grid search for *Q* ∈ [1, *M*] and retain the value of Q which maximizes the likelihood function as shown below:

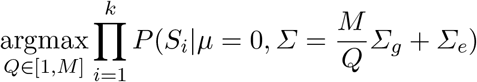

## A UK Biobank Data

We used four traits from the UK Biobank released in 2017 (and in 2018) as the basis of our simulations [Neale Lab, 2017, 2018]. For both sets of summary statistics, only the variants which are biallelic, have non-ambiguous strands, a minor allele frequency greater than 1%, an INFO score greater than 80%, and found in the 1000 Genomes European reference panel are retained [Consortium, 2015]. We used LD-Score regression [Bulik-Sullivan et al., 2015a] to calculate the genetic variance of each trait as shown in Table 6 and Table 8. For calculating genetic covariance and environmental covariance, we used cross-trait LD-Score [Bulik-Sullivan et al., 2015b]. The genetic covariance produced by the software is reported in Table 7 and Table 9. By taking the intercept and rescaling it by the 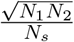 where *N*_1_ is the sample size in trait 1, *N*_2_ is the sample size in trait 2, and *N_s_* is the number of overlapping individuals, we are able to recover the phenotypic covariance. By subtracting the genetic covariance from the phenotypic covariance, the environmental covariance is estimated. As we use summary statistics and the true sample overlap is unknown, we assume no trait specific missing individuals, and we, therefore, set *N_s_* = *min*{*N*_1_, *N*_2_}.

**Table 6:** The genetic variance and sample size from 2017 release used in simulations. We used summary statistic for body mass index, diastolic blood pressure, height, and systolic blood pressure from the 2017 release of the UK Biobank as input. We reported the sample size, genetic variance estimated by LD-Score regression and the LD-Score intercept.

| Trait | Genetic Variance | Sample Size | LD-Score Intercept |
| --- | --- | --- | --- |
| body mass index ($\mathcal{B}$) | 0.241 | 336,107 | 1.082 |
| diastolic blood pressure ($\mathcal{D}$) | 0.129 | 317,756 | 1.074 |
| height ($\mathcal{H}$) | 0.429 | 336,574 | 1.237 |
| systolic blood pressure ($\mathcal{S}$) | 0.114 | 317,754 | 1.108 |

**Table 7:** The genetic and environmental correlation used in simulations from 2017 version of UK Biobank data. We used summary statistics for body mass index, diastolic blood pressure, height, and systolic blood pressure from the 2017 release of the UK Biobank as input. We use cross-trait LD-Score regression to estimate the genetic and environmental correlation and report the LD-Score intercept.

| Trait 1 | Trait 2 | Genetic Correlation | Environmental Correlation | LD-Score Intercept |
| --- | --- | --- | --- | --- |
| body mass index ($\mathcal{B}$) | diastolic blood pressure ($\mathcal{D}$) | 0.305 | 0.258 | 0.303 |
| body mass index ($\mathcal{B}$) | height ($\mathcal{H}$) | -0.165 | -0.084 | -0.137 |
| body mass index ($\mathcal{B}$) | systolic blood pressure ($\mathcal{S}$) | 0.166 | 0.198 | 0.219 |
| diastolic blood pressure ($\mathcal{D}$) | height ($\mathcal{H}$) | -0.125 | -0.004 | -0.025 |
| diastolic blood pressure ($\mathcal{D}$) | systolic blood pressure ($\mathcal{S}$) | 0.648 | 0.686 | 0.765 |
| height ($\mathcal{H}$) | systolic blood pressure ($\mathcal{S}$) | -0.144 | -0.104 | -0.132 |

**Table 8:** The genetic variance and sample size used in simulations and real data analysis. Using the summary statistics for body mass index, diastolic blood pressure, height, and systolic blood pressure from the 2018 release of the UK Biobank, we estimate the genetic variance with LD-Score regression. We report the sample sizes, genetic variance, and LD-Score intercept.

| Trait | Genetic Variance | Sample Size | LD-Score Intercept |
| --- | --- | --- | --- |
| body mass index ($\mathcal{B}$) | 0.243 | 359,983 | 1.057 |
| diastolic blood pressure ($\mathcal{D}$) | 0.132 | 340,162 | 1.068 |
| height ($\mathcal{H}$) | 0.469 | 360,388 | 1.270 |
| systolic blood pressure ($\mathcal{S}$) | 0.139 | 340,159 | 1.070 |

**Table 9:**
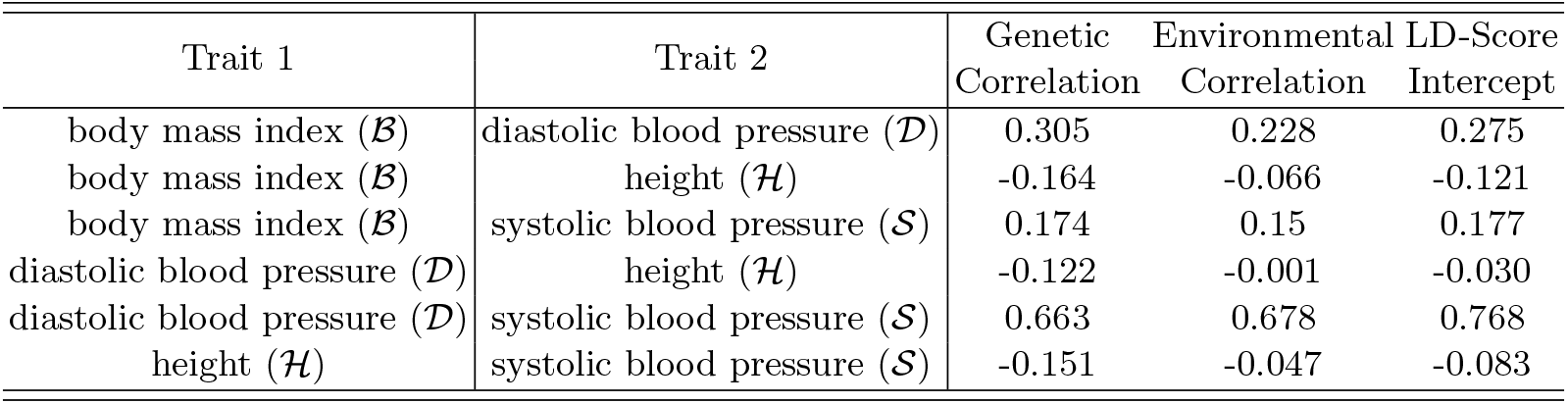
The genetic and environmental correlation used in real data analysis and simulations. We used summary statistics for body mass index, diastolic blood pressure, height, and systolic blood pressure from the 2018 release of the UK Biobank as input to cross-trait LD-Score regression. We report the genetic and environmental correlation as well as the LD-Score intercept.

| Trait 1 | Trait 2 | Genetic Correlation | Environmental Correlation | LD-Score Intercept |
| --- | --- | --- | --- | --- |
| body mass index ($\mathcal{B}$) | diastolic blood pressure ($\mathcal{D}$) | 0.305 | 0.228 | 0.275 |
| body mass index ($\mathcal{B}$) | height ($\mathcal{H}$) | -0.164 | -0.066 | -0.121 |
| body mass index ($\mathcal{B}$) | systolic blood pressure ($\mathcal{S}$) | 0.174 | 0.15 | 0.177 |
| diastolic blood pressure ($\mathcal{D}$) | height ($\mathcal{H}$) | -0.122 | -0.001 | -0.030 |
| diastolic blood pressure ($\mathcal{D}$) | systolic blood pressure ($\mathcal{S}$) | 0.663 | 0.678 | 0.768 |
| height ($\mathcal{H}$) | systolic blood pressure ($\mathcal{S}$) | -0.151 | -0.047 | -0.083 |

For simulations, PAT will use the reported values to define its likelihood ratio test; MI GWAS will, however, will make no assumption about the phenotypic covariance. HIPO and MTAG define their parameters slightly differently, but all methods use the same results from LD-Score regression and cross-trait LD-Score regression. For simulations comparing the power between MI GWAS and PAT, we set the environmental correlation to 0% for all traits. Simulations comparing PAT to MI GWAS and for showing the stability of null simulations use the 2017 version of the UK Biobank summary statistics (see Table 6 and Table 7 for values). All other simulations and real data analysis use the summary statistics from 2018 (see Table 8 and Table 9). The switch in summary statistic version is due to the 2017 version of the UK Biobank results being no longer available which prevents reproduction of results.

## B Omnibus Associations in the UK Biobank

In this real data analysis, we analyze summary statistics for body mass index 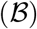, diastolic blood pressure 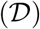, height 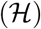, and systolic blood pressure 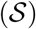 measured in the UK Biobank. Here, five methods are compared: Single Trait GWAS (how the z-scores and p-values were derived), HIPO, MTAG, MI GWAS, and PAT [Qi and Chatterjee, 2018, Turley et al., 2018]. There are 7,025,734 variants which are biallelic, have non-ambiguous strands, a minor allele frequency greater than 1%, and an INFO score greater than 80%. The reference and alternate allele are coordinated across traits by flipping the direction of the effect when necessary. LD-Score regression and cross-trait LD-Score regression are used to calculate the genetic and environmental covariance structure (see above) [Bulik-Sullivan et al., 2015a,b].

The first column lists all subsets of the four traits while the second column contains the results from Single Trait GWAS. This is the maximum number of variants the other four methods can recapture. Each row is based on which traits are identified as associated using individual level data for each trait separately. For example, the first row shows the results for body mass index 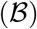, diastolic blood pressure 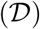, height 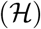, and systolic blood pressure 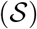. There are 176 unique variants found to be significantly associated in all four traits by Single Trait GWAS; therefore, HIPO MTAG, MI GWAS, and PAT cannot have a value greater than 176 in the first row. In Table 10, we see every method except HIPO is able to identify all 176 variants as associated with at least one of the four traits. We note that while MTAG is not an omnibus method, a variant is deemed associated as long as one trait is significantly associated. Additionally, HIPO computes four components for testing. If at least one component is genome-wide significant, the variant is interpreted as associated. All methods are tested at *α* = 5 × 10^−8^ and bound by the original single trait results to provide a fair comparison to fundamentally different methods.

**Table 10:**
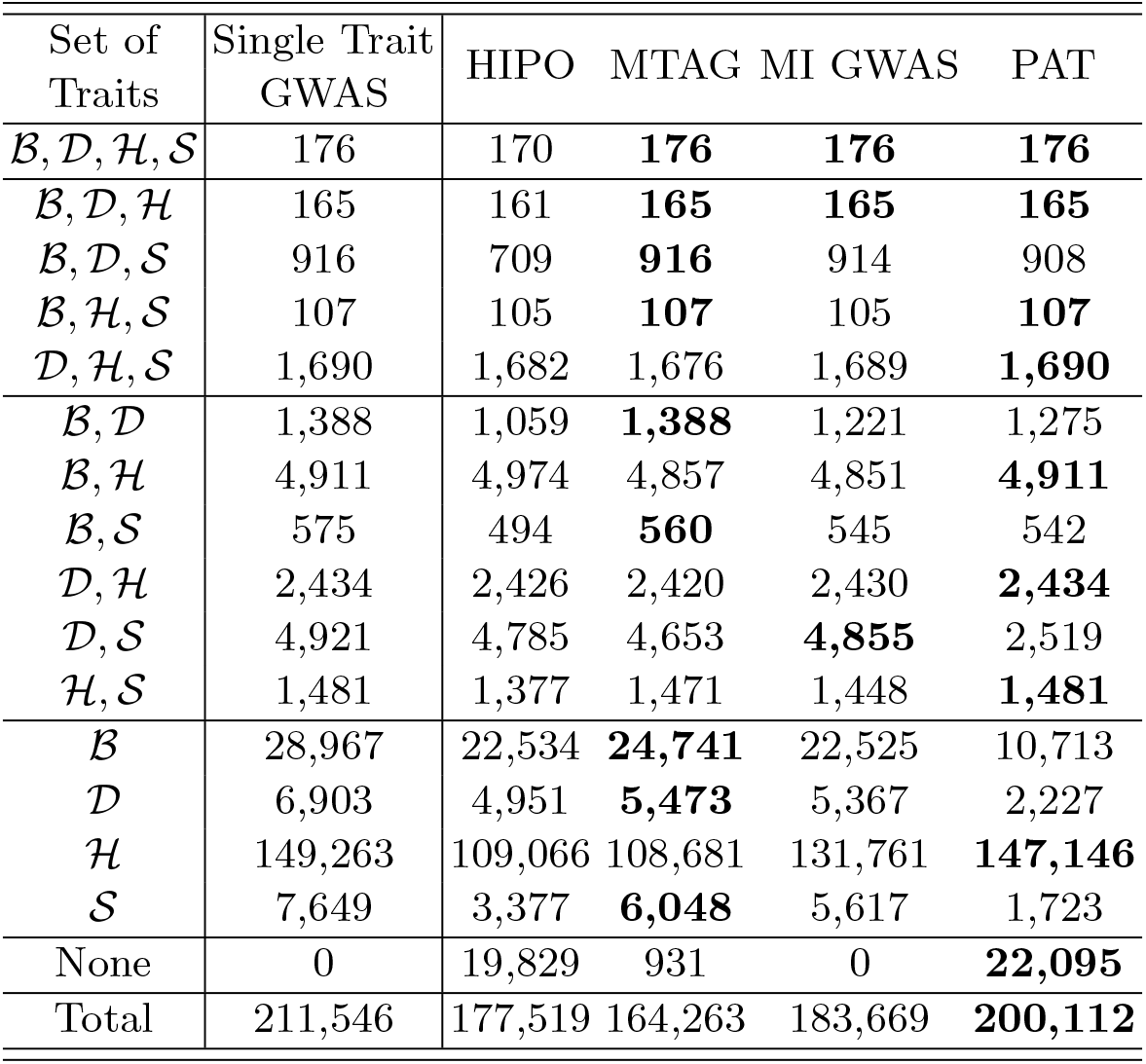
UK Biobank data analysis. The first column lists the set of traits. Each remaining column corresponds to different GWAS methods with the second column showing the Single Trait GWAS results. These results were generated from individual level data and produced the summary statistics. The remaining columns show how many of these variants were also identified by HIPO, MTAG, MI GWAS, and PAT, respectively. We separate the data according to Single Trait GWAS significant results; for example, the first row of results is the number of variants found associated with all traits 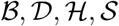 (body mass index, diastolic blood pressure, height, and systolic blood pressure).

| Set of Traits | Single Trait GWAS | HIPO | MTAG | MI GWAS | PAT |
| --- | --- | --- | --- | --- | --- |
| $\mathcal{B}, \mathcal{D}, \mathcal{H}, \mathcal{S}$ | 176 | 170 | **176** | **176** | **176** |
| $\mathcal{B}, \mathcal{D}, \mathcal{H}$ | 165 | 161 | **165** | **165** | **165** |
| $\mathcal{B}, \mathcal{D}, \mathcal{S}$ | 916 | 709 | **916** | 914 | 908 |
| $\mathcal{B}, \mathcal{H}, \mathcal{S}$ | 107 | 105 | **107** | 105 | **107** |
| $\mathcal{D}, \mathcal{H}, \mathcal{S}$ | 1,690 | 1,682 | 1,676 | 1,689 | **1,690** |
| $\mathcal{B}, \mathcal{D}$ | 1,388 | 1,059 | **1,388** | 1,221 | 1,275 |
| $\mathcal{B}, \mathcal{H}$ | 4,911 | 4,974 | 4,857 | 4,851 | **4,911** |
| $\mathcal{B}, \mathcal{S}$ | 575 | 494 | **560** | 545 | 542 |
| $\mathcal{D}, \mathcal{H}$ | 2,434 | 2,426 | 2,420 | 2,430 | **2,434** |
| $\mathcal{D}, \mathcal{S}$ | 4,921 | 4,785 | 4,653 | **4,855** | 2,519 |
| $\mathcal{H}, \mathcal{S}$ | 1,481 | 1,377 | 1,471 | 1,448 | **1,481** |
| $\mathcal{B}$ | 28,967 | 22,534 | **24,741** | 22,525 | 10,713 |
| $\mathcal{D}$ | 6,903 | 4,951 | **5,473** | 5,367 | 2,227 |
| $\mathcal{H}$ | 149,263 | 109,066 | 108,681 | 131,761 | **147,146** |
| $\mathcal{S}$ | 7,649 | 3,377 | **6,048** | 5,617 | 1,723 |
| None | 0 | 19,829 | 931 | 0 | **22,095** |
| Total | 211,546 | 177,519 | 164,263 | 183,669 | **200,112** |

We now consider the case where the variant is associated with 3 of the traits for single trait GWAS. Under this condition, all of the summary statistic methods correctly identify most associated variants. HIPO is the only method that fails to recover all associated variants under any of the scenarios of an effect in 3 of the 4 traits though it recovers the vast majority. MTAG recovers all associated variants for 3 of the 4 sets, but it missed 14 variants when considering 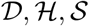. PAT also successfully returns all associated variants for 3 of the 4 sets, but missed 8 variants when considering 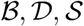. MI GWAS missed 1 or 2 variants in 3 of the 4 sets of traits but overall recovered most variants.

For variant originally found to be associated with 2 of the 4 traits, the various summary statistic methods still perform similarly. The major exception is for diastolic blood pressure 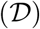 and systolic blood pressure 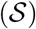 where PAT only identifies 2,519 of the 4,921 variants while HIPO, MTAG and MI GWAS found 4,785, 4,653, and 4,855, respectively. As 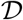 and 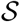 are highly genetically and environmentally correlated, it is not surprising that PAT recovers fewer variants. This is because PAT is more conservative when summary statistics are genetically and environmentally correlated in the same direction to control false positives (see Fig. 2). When a variant is originally associated with only one trait, we find general PAT generally identifies significantly fewer variants than HIPO, MTAG, and MI GWAS. The exception is height 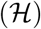. This is likely due to the other traits being more genetically correlated than they are environmentally correlated to 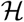. This along with the fact that there are many large peaks in 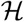 could result in PAT identifying more variants than the the other methods.

The second to last row is the number of variants identified as novel associations not originally found by the Single Trait GWAS. Here, MI GWAS finds no new associations, but this is by design. MTAG, however, is able to identify 931 additional variants. HIPO identifies 19,829 while PAT discovers 22,095.

In the results shown here, HIPO fails to to recapture as many associations discovered using single trait GWAS as the other three methods. MTAG, MI GWAS and PAT, however, performed more similarly to each other. When it comes to identifying novel associations, HIPO and PAT are far more powerful than MTAG and MI GWAS. Overall, 37,890 novel associations were discovered by three of the multi-trait methods (HIPO, MTAG, and PAT). 385 of these variants are found by all three methods. 17,788 out of the 22,095 variants discovered by PAT are unique to PAT. For HIPO, 15,407 out of its 19,829 variants are unique while 115 of MTAG’s 931 associations are only identified by MTAG. While PAT is the most powerful in regards to identifying novel associations, these results indicate HIPO, MTAG, and PAT are able to identify many unique variants due to their differing model designs.

## C Empirical M-value Threshold

Through simulations, we show that the m-value framework has a low false positive assignment rate. M-values, however, cannot be calibrated the same way p-values can be by using significance thresholds. Simulations both in the original m-value paper [Han and Eskin, 2012] and here support the use of 0.90 as the threshold for interpretation. This is to say, if the m-value is greater than 0.90, the user should interpret the variant as associated with the trait.

While a reasonable threshold, the user will need to prioritize variants for further exploration and follow up. Here, we explore the true positive rate of the m-value interpretation in ranked order. In results in the main paper, we present two tables, Table 1 and Table 2 from 700,000 simulated variants and their z-scores for four traits. 70,000 (10%) of the variants have a genetic effect in at least one trait which were evenly split across seven configurations of genetic effect. The list of configuration of genetic effect are stated below:

1. genetic effect in body mass index, diastolic blood pressure, height and systolic blood pressure 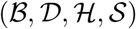
2. genetic effect in body mass index, diastolic blood pressure and height 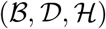
3. genetic effect in diastolic blood pressure, height, and systolic blood pressure 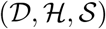
4. genetic effect in body mass index, and height 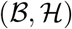
5. genetic effect in diastolic blood pressure and systolic blood pressure 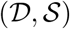
6. genetic effect in body mass index 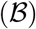
7. genetic effect in height 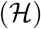

The first step is to use PAT (or HIPO) to assign omnibus p-values to the variants [Qi and Chatterjee, 2018]. While PAT only performs one test, HIPO computes 4 weighting components for testing. A variant is considered associated if found genome-wide significant for at least one component. For both methods, the associated variants are assigned an m-value for each trait. For each per trait interpretation, the corresponding ground truth is known. If the variant genetically affects a trait and the m-value is greater than 0.9, we consider this a true positive. If the m-value is greater than 0.9 and the variant does not have a genetic effect, this is a false positive. We define true and false negatives using the same logic for m-values less than or equal to 0.9.

In Table 1, we record that PAT identifies 3,569 associated variants while HIPO identifies 2,496 associated variants. Therefore, the respective total number of per trait m-values across four traits is 14,276 and 9,984. The m-values for each method are ranked from largest to smallest [1.0, 0.0] and binned in sets of 100. For each bin, we calculate the true positive rate and show the results in Fig. 8. In addition to the true positive rate for each bin, we also indicate the minimum m-value in each bin. In Fig. 8, The x-axis denotes bins of m-values in ranked order while the y-axis is the true positive rate.

**Fig. 8:**
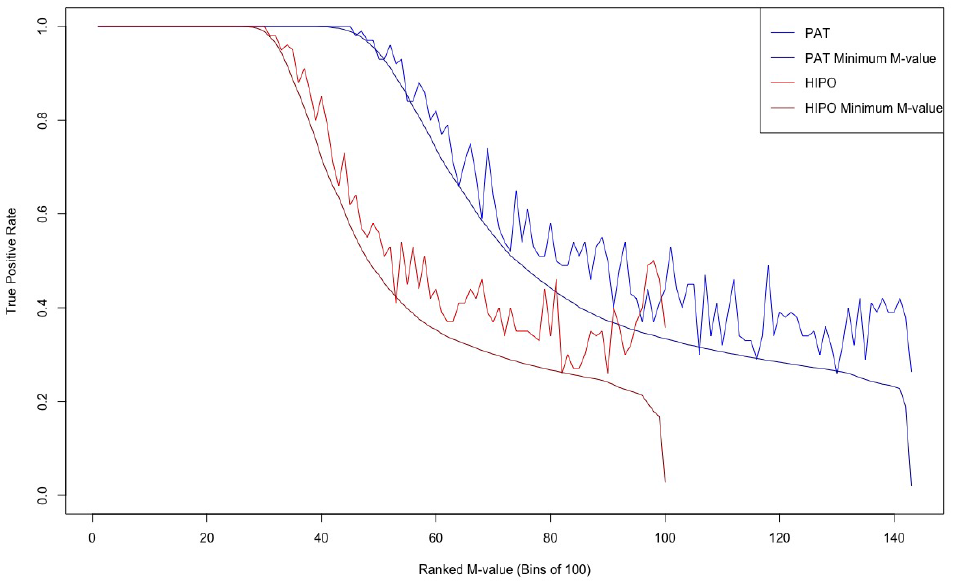
Empirical M-value Threshold. The m-value framework is applied to variants identified as associated using PAT and HIPO. The resulting m-values are ranked and binned in sets of 100. For each bin, we plot the smallest m-value as well as the true positive assignment rate for both methods.

In Fig. 8, we see that true positive rate and the minimum m-value for each bin follow the same general pattern. This means that each subsequent set of per trait associations, the proportion of true positives can be coarsely estimated by the set’s minimum m-value. While this approach does not provide the false positive rate for any particular m-value, it may provide some guidance on whether a particular subset of variants has a high enough true positive rate to warrant further analysis.

To make this explanation more concrete, the true positive rate for PAT in the first 45 bins (4,500 m-values) is 100% for each. Using three digits of precision, the first 39 bins have a minimum m-value of 1.000. While bins 40-45 also have a true positive rate of 100%, the minimum m-value observed for those bins is less than 1.000 with the minimum m-value for bin 45 being 0.993. While we know the ground truth for the 45th bin that there are 0 false positive, we could also use the minimum m-value to estimate that approximately 1 of the m-values is a false positive. This approach can be used as a rough lower bound when determining whether further exploration of a particular set of variants is warranted. From this, a rough estimate for all m-values greater than 0.9 is a 90% true positive rate. This is far more conservative than necessary as we empirically have shown false positive rates of 0.6% or less. Instead, by using smaller bins (such of size 100) the user can have a crude estimate of the true positive rate of that bin of m-values.

## D Ranked false positive rate between m-values and p-values

In the main paper using simulations, we compare the per trait m-values produced for PAT and HIPO to the per trait p-values produced by MTAG. In Table 2, PAT identifies 5,223 true per trait associations. Applying m-values results in 3,430 per trait associations for HIPO while MTAG discovers 2,444. While these results indicate PAT is the most powerful approach, they also provide evidence that in general computing posterior predictions (m-values) after omnibus associations is a more powerful approach to association testing than directly analyzing each trait using MTAG.

While the number of true positives support this claim, the difference in the false positive rate between m-values and p-values may draw this claim into question. This is to say, if the same number of false positives were produced by m-values and MTAG’s p-values, it is possible that MTAG would be the most powerful approach.

In order to test this claim, m-values must be modified to better reflect p-values. Currently, m-values are only interpreted for variants deemed genome-wide significant. This is due to their design as an interpretation framework. M-values are not designed to replace p-values but to elucidate what traits may be driving the significant association. In this comparison, we assign an m-value to every z-score instead of only to the omnibus significant z-scores. This enables a comparison of all posterior predictions to the per trait p-values of MTAG. While illustrative of the power of m-values, this approach is not advisable in practice. M-values should only be used as a means of interpreting omnibus associations.

For this comparison, we will use the 700,000 simulations with 10% causal variants previously described above. The 70,000 causal variants will be equally split across seven configurations of genetic effect. Under these seven configurations, the traits genetically affected by the variant will vary. For each of the configurations, three different effect sizes will also be modeled. As stated previously, MTAG directly produces a per trait p-value and m-values are assigned to HIPO and PAT.

In Fig. 9, we rank the m-values for PAT and HIPO and the p-values for MTAG. For p-values, the rank order is from [0.0,1.0] while m-values is ordered from [1.0, 0.0]. For each m-value (and p-value), whether or not that the variant is truly associated with the trait is known; therefore, the true positive rate for the variants in rank order can be calculated. After ranking the m-values (and p-values), the variants are binned in sets of 1,000 for each method. In Fig 9, sub-figure (A), the first 200 bins or 200,000 top p-values (m-values) for each method are along the x-axis. Along the y-axis, the true positive rate for each bin is shown. For the most significant associations by p-values and m-values the true positive rate is 1. As the ranked position decreases, the true positive rate decreases. Overall, the general trend of m-values for both HIPO and PAT and p-values for MTAG follow the same pattern. In the top left of sub-figure (A) MTAG (purple) can be seen as having a slightly lower true positive rate than PAT (blue) and HIPO (red) which overlay each other.

**Fig. 9:**
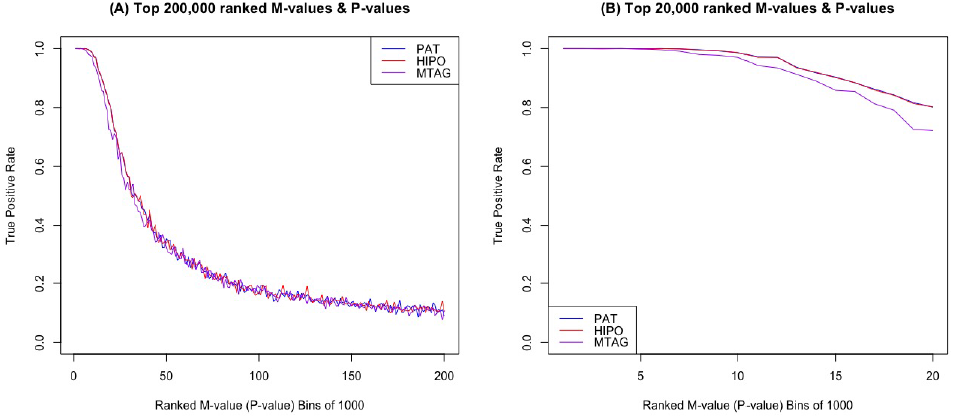
Comparison of m-values and p-values. M-values are assigned to all z-scores for PAT and HIPO. For each method, they are ranked and placed in bins of 1,000. The p-values from MTAG are also ranked and binned in sets of 1,000. A comparison of their respective true positives rates are shown in (A) the first 200 bins and (B) first 20 bins.

In Fig. 9 sub-figure (B), we explore the top 20 bins or 20,000 m-values and p-values more closely. Here, we see that there is some separation between the m-values for PAT and HIPO, respectively shown in blue and red, and MTAG in purple. From this, there is evidence that while the m-value framework does not control for false positives directly, there is a gain in power relative to a directly interpretable multi-trait method, such as MTAG. We note that while true, neither the true (nor false) positive rate can be elucidated from the m-value directly. Therefore, m-values should only be used as designed to interpret significant p-values.

